# Cross immunity protection and antibody dependent enhancement: A distributed delay dynamic model

**DOI:** 10.1101/2021.07.22.453430

**Authors:** Vanessa Steindorf, Sergio Oliva, Jianhong Wu

## Abstract

Dengue fever is endemic in tropical and sub-tropical countries, and some of the important features of Dengue fever spread continues to pose challenges for mathematical modelling. Here, we propose a system of integro-differential equations (IDE) to study the disease transmission dynamics that involves multiserotypes and cross immunity. Our main objective is to incorporate and analyze the effect of a general time delay term describing acquired cross immunity protection and the effect of antibody dependent enhancement (ADE), both characteristics of Dengue fever. We perform qualitative analysis of the model and obtain results to show the stability of the epidemiologically important steady solutions that is completely determined by the basic reproduction number and the invasion reproduction number. We establish the global dynamics, by constructing suitable Lyapunov functions. We also conduct some numerical experiments to illustrate bifurcation structures, indicating the occurrence of periodic oscillations for specific range of values of a key parameter representing the ADE.

## 1 Introduction

Dengue fever is an endemic disease in tropical and sub-tropical countries. According to the World Health Organization WHO (2018), in 2016, more than 2.38 million cases were reported in the Americas, where Brazil alone contributed with almost 1.5 million cases. In 2017, a significant reduction in the number of cases in Americas was reported, even though a recent estimate indicates 390 million cases per year around the world (WHO, 2018).

Dengue fever is caused by Dengue virus (DENV), which is transmitted to a human population by the bite of infected mosquitos. It is known that there are four distinct virus serotypes circulating in the population (Gubler, 2014). The disease presents as Dengue classic or Dengue hemorrhagic fever (DHF).

Antibody-Dependent Enhancement (ADE) was proposed to explain the more frequent occurrence of severity of Dengue fever and Dengue hemorrhagic fever in secondary infections (Guzman, 2013; Gubler, 2014; Reich, 2013). Another characteristic of Dengue virus is the cross immunity protection, so that the infection with any of the four serotypes leads to a short term protection for all serotypes. Long term protection occurs only for the strain which the individual was infected (Reich, 2013).

Some mathematical models that include ADE were proposed by Adams (2006), Bianco (2009), Billings (2007) and Hu (2013). Adams (2006) included in a two-strain model the ADE effect and, described the immunological distance between dengue serotypes through a function with the hypotheses that this function reduces the probability of contracting a secondary infection. Numerical results were presented in that study.

Aguiar (2007) proposed an Ordinary Differential Equations (ODE) system for two serotypes of Dengue virus. The total population was stratified by their clinical states. Time series simulations was investigated, and various bifurcation phenomena were observed. Theoretical mathematical analysis of the model of Aguiar (2007) was made by Aguiar (2008) and Kooi (2014). Aguiar (2008), quantified the attractor structure, limit cycle and chaotic attractor by calculating Lyapunov exponents. Analytic formulations for the equilibrium and analysis of the bifurcation structure were obtained by Kooi (2014) for the symmetric case (when the strains are the same).

These models aforementioned are formulated in terms of ordinary differential equations, without incorporating time lags for recovery and incubation. General delay models was introduced by Van den Driesshe (1996), using a special step function for the recovery period. Recently, Nah (2014) described a mathematical model for Malaria transmission with a time delay involved during the exposure and a general incubation period.

A mathematical model describing a general multi-strain disease was proposed by Chen (2016). The authors proposed a delay diffusive two strain disease model, considering the SIR structure, constant recruitment rate and, constant time delay representing the length of immunity period. The stability of the model was determinate by the Basic Reproduction Number. Guan (2017) also described a Dengue fever model with time delay. The time delay included in the model refers to a time incubation of the virus in an infected population and in an infected mosquito population.

Taking into account a simpler SIR model, Huang (2016) considered an infinite distributed delay on complex population network. Numerical experiments confirmed that the delay slows down the extinction of the disease when the threshold value is smaller than 1, while the delay accelerates the spreading if it is bigger than 1. Distributed delay was also used by Xu (2017) in a SVEIR model involving a continuous vaccination strategy, his results showed that distributed delay had no impact on the qualitative behaviour and global dynamics.

Since Dengue fever constitutes a public health problem, we proposed a mathematical model, namely, an IDE system that can be applied to describe and to study the propagation of Dengue fever in a population, considering two main characteristics of the disease: ADE and cross immunity protection. The main purpose is to include and analyze the effect of the general time delay on the model, included in order to represent the length of the cross immunity protection. In addition, a constant parameter will be added in the model in order to represent the effect of ADE.

In the third section, the model is analyzed. Defining the associated limiting system, we look for equilibriums and show local stability that is determined by an important threshold value. At section four, by constructing Lyapunov functions we prove global stability. Numerical analysis are made at section five, where the bifurcation structure and stability of the coexistence equilibrium are numerically studied. Final considerations and conclusions are made at last section.

## 2 Model Formulation

In this section we are going to propose a model to describe a multi-strain disease, motivated by Dengue fever.

Let *N* (*t*) be the total population of individuals at time *t* in a region. We divided the population in disjoint classes according to the individual status, susceptible for all serotypes, infected by serotype *i*, temporarily immune for all serotypes after being infected by serotype *i*, recovered for serotype *i* but susceptible to the others and, recovered for all serotypes, represented at time *t*, respectively by *S*(*t*), *I*_*i*_(*t*), *C*_*i*_(*t*), *R*_*i*_(*t*) e *R*(*t*). Furthermore, we include two more classes for the population, *I*_*ij*_(*t*), representing the sub-population reinfected by serotype *j* after being infected by serotype *i*, with *i, j* = 1, 2.

Considering that birth rate and mortality rate are equal, let *d* be a constant natural mortality rate for human population. We assume that the constant rate *β*_*i*_ is the transmission rate by serotype *i*, for the first time that an individual is infected. Whereas the constant rate *α*_*j*_ is the transmission rate, by different serotype, by serotype *j*, for the second time that an individual is infected.

Note that the mosquito population is not considered explicitly in the model, the rate *β*_*i*_ and *α*_*j*_ will be the average number of bite and, the probability of a susceptible individual being bite for an infected mosquito by serotype *i* and, the probability of a recovered individual by serotype *i* being bite again for an infected mosquito by serotype *j*, respectively.

Individuals in the infectious classes *I*_*i*_(*t*), remain in this class with average time 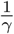, since we assume that the length in this class is exponentially distributed. Once in infected class, the individual recovers and goes to the temporary immunity class *C*_*i*_(*t*). In this class, the individual gets temporary immunity for all serotypes.

Moreover, we assume for *C*_*i*_ class a general length of immunity. Let *P*_*i*_(*t*) be the fraction of individuals that after recovered from the serotype *i*, they remain cross protected by all serotypes, *t* units after get in the temporary immunity class. It is reasonable to assume that

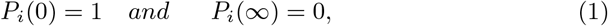

and, satisfies that *P*_*i*_(*t*) is non-increasing and

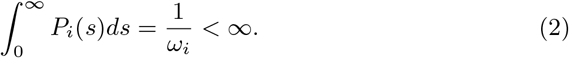

Thus, the number of cross protected individuals is given by

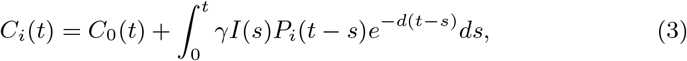

where *C*_0_ is the number of protected individuals at time *t* = 0 that are still immune at time *t*. Of course *C*_0_ must satisfy lim_*t*→∞_ *C*_0_(*t*) = 0.

After the cross protection time, the individual remains susceptible for the other strains. In this way, an individual in the *R*_*i*_ class can be infected again with a rate *α*_*j*_ by a different serotype. Only after being infected by all strains the individual became immune, recovering with constant rate *γ*, remaining life long recovered.

Whenever the individual is infected again, by a different serotype, the immune system responds increasing the infectiousness and enhancing viral replication (Gubler, 2014; Guzman, 2013; Katzelnick, 2017). This effect is called ADE, when secondary infections are possible in the presence of intermediate level of antibody, that make growths the viral replication (Gubler, 2014; Guzman, 2013; Katzelnick, 2017).

Adams (2006), Bianco (2009),Hu (2013),Aguiar (2007), Aguiar (2008), Kooi (2014) and Billings (2007) assume that this increasing viral load is associated with increasing transmission infection, and they use this assumption to describe mathematically and to model ADE using a constant *ϕ* that represents the increase of the transmission rate, associated to the transmissibility of the recovery individuals transmit infections.

According to Katzelnick (2017), ADE occurs at a specific range of antibody concentrations. Low levels of antibody did not enhance disease, intermediate levels exacerbated disease, and high levels protected against severe disease. The authors verify enhancement in humans and show that pre-existing antibodies is associated with the severity of the disease, and they also showed that the immune correlate for enhanced severe dengue is distinct from that for protection. Futhermore, Rohman (2011) affirms that depending on the specific antibody concentration, dengue virus antibodies can inhibit viral infection (neutralization) or enhance infection.

Therefore, ADE happens or not, increasing or decreasing infection, due to high or low antibodies concentration on people who was infected once (recovered) and again infected. Clearly, this high viral load is not related to the increasing transmissibility (transmission from recovered individual to new infections) but, rather to the capacity of immune response system of primary individual be able to respond to the secondary infection, and then, to be protected against infection or enhance the disease.

Thus, the ADE characteristic must be described as a constant rate that can increase or decrease the probability of a recovery individual, already infected once, be infected again, assuming that the individual with primary infection carries viral load that allows to neutralize or enhance infection. In this way, in this study, we are proposing that *Φ* represents the fraction that decreases (*ϕ* < 1) or increases (enhance) (*ϕ* > 1) the probability of secondary infections in primary infected individuals. This is, the epidemiological consequence ADE will be described through the constant coefficient *ϕ*, where it is assumed that previously exposure to one serotype results in increasing of the susceptibility for reinfection.

Thus, based on the assumptions and in the studies by Wang (2016), Cooke (1996), Van den Driesshe (2007), and taking the derivative of equation (3), assuming *C*_0_ = 0, the model can be described as follows:

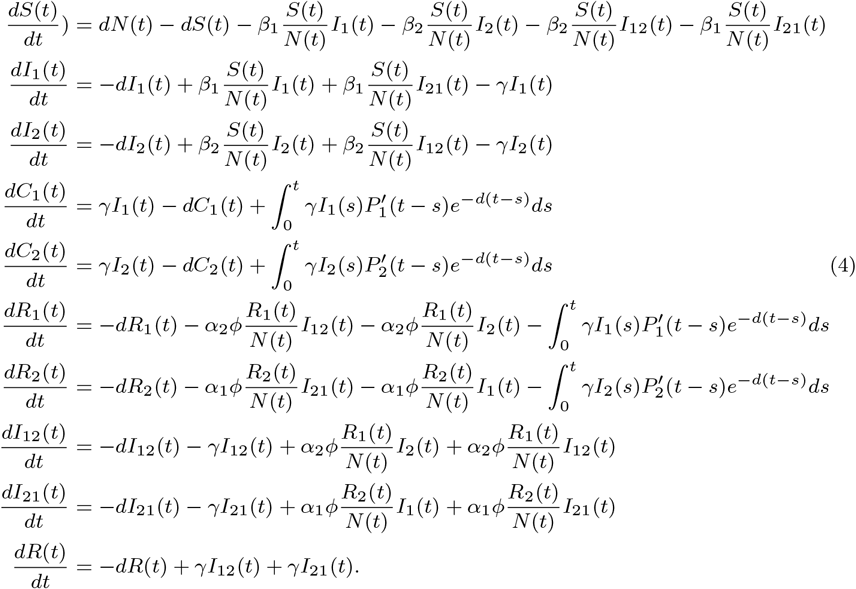

The total population dynamics is determined by

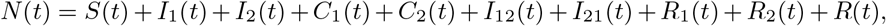

and, since 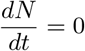, the total population remains constant in time.

Denote 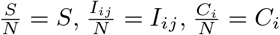 and 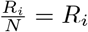 representing, for each class, the fractions of the population, and *N* = *N* ^*\**^. Thus, the sum of the total population satisfies *S* + *I*_1_ + *I*_2_ + *C*_1_ + *C*_2_ + *R*_1_ + *R*_2_ + *I*_12_ + *I*_21_ + *R* = 1. In addition, the dynamics for the recovered and cross immunity classes are decoupled. Therefore, the original system can be studied by analyzing the following subsystem:

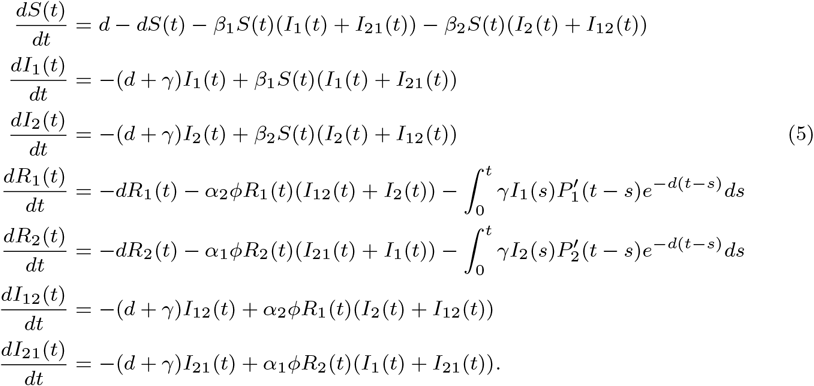

This subsystem will be qualitatively analyzed at section (3). In the next sub-section, we are going to discuss a particular case of this model.

### 2.1 Particular case: Exponential immunity

In this, subsection we are going to discuss a particular case of the model (4). If we assume the length of immunity being exponentially distributed, which means that the fraction individuals temporarily immune remain in the *C*_*i*_ class is 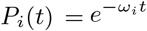, with *ω*_*i*_ > 0, for *i* = 1, 2, then the IDE system (4) became an ODE system. Thereby, the system already normalized with the variables representing the fractions of the populations, can be described as follows:

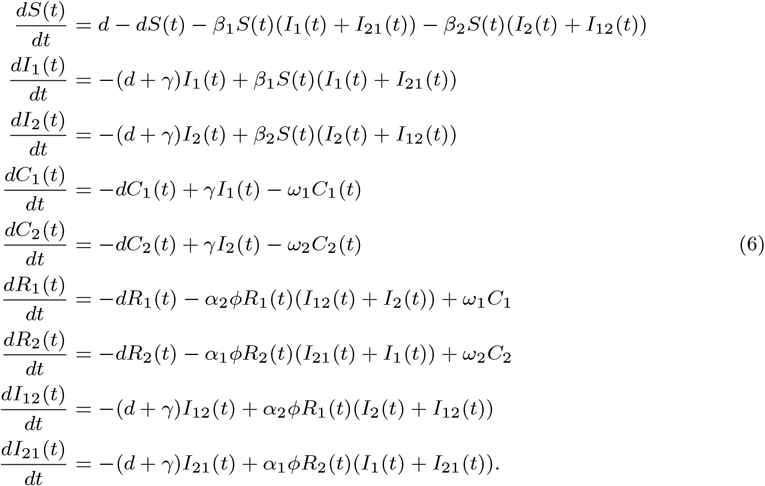

Qualitatively analysis was made for this particular model through classic theory for ODE systems. Details are found in the supplementary material. We summarize the results here, but first, we are going to define a threshold value that is very important for the stability and existence of equilibriums.

#### 2.1.1 Basic Reproduction number and Invasion Reproduction number

The Basic Reproduction number ℛ_0_ is defined for many authors, as by Van den Driesshe (2008), as the expect number of secondary infections produced by one case in a susceptible population and, also as a measure of the potential for disease spread in a population. Mathematically, the Basic Reproduction Number is a threshold for stability of Disease Free equilibrium (Van den Driesshe, 2008).

We will define 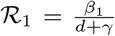 as the Basic Reproduction Number for strain one. And, 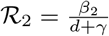 as the Basic Reproduction Number for strain two.

It is usual to define an overall Reproduction Number for a multi-strain model, with different strains. Thus, it will be defined as:

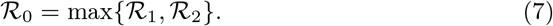

This definition, according to Martcheva (2015), is similar with that given by Van den Driesshe (2008), where the Basic Reproduction number is defined mathematically as the spectral radius (the maximum of the modulus of the eigenvalues) of the Next Generation matrix.

It is usual to find another important threshold value when working with multiple strain model. This value will determine whether the Coexistence Endemic equilibrium (CEE) will be in the biological positive region and, also the stability of the Boundary Equilibrium (BE). This threshold is called Invasion Reproduction number (of strain one at the equilibrium of strain two) (Martcheva, 2015), and we are going to denote by ℛ_*Inv*_. Martcheva (2015) defined epidemiologically as a number of secondary infections that one individual infected with the strain one will produce in a population, in which the strain two is at equilibrium during its infectious lifetime.

### 2.2 Discussion of Qualitative Result

For the ODE model (6) we proved the existence of four equilibriums (proofs and details can be found in the supplementary material). The Disease Free equilibrium (DFE) is locally asymptotically stable if ℛ_0_ < 1, and thus, the disease will die out. While, the DFE is unstable if ℛ_0_ > 1.

We found two BE, each equilibrium corresponding to only one infection. The infection with the smallest ℛ_*i*_ is always unstable, while the other is stable if ℛ_*Inv*_ < 1. This means that the disease with the biggest force will persist and the other will die out. In addition, the BE is unstable if ℛ_*Inv*_ > 1 and, thus, there is a forth equilibrium, the CEE, in the positive invariant region. In this scenario, the two strains will coexist.

This ODE model is very similar to the models proposed by Aguiar (2007),Aguiar (2008),Kooi (2014). However the assumptions for the ADE effect is different, leading to different dynamics. There is no mention of the CEE in the works cited and studied by Aguiar (2007, 2008), Kooi (2014). The analytical result for the CEE and the stability was obtained only for the symmetric case by Aguiar (2008), when all the parameters are equal for different strains.

We could proved that for certain values of the parameter that represents the enhancement, there is indeed a CEE, within the region. Which is a very important result because, according to VinodKumar (2013), different serotypes have been coo-circulating in the same area with one of them being dominant during an outbreak.

Also, we proved numerically that only for small parameter values that represent the ADE the equilibrium can be stable. Additionally, we showed that bifurcation occurs and, for the most of the values of the parameter *ϕ*, that represent the ADE, the equilibrium exist and it is unstable. Thus, the coexistence of the two serotypes is possible but not established and the solution oscillates for a critical parameter value.

## 3 General Case: Time delay system

In the previous section (2.2) we discuss a particular case, when the time of cross-immunity protection is defined exponentially distributed. We will now to analyze the model when the cross-immunity protection time can be defined as any continuous function with some additional properties.

Consider the system (5) together with the assumptions (1) and (2). Following the idea in (Brauer, 1978), we will examine the system (5) as a perturbation of the limiting system:

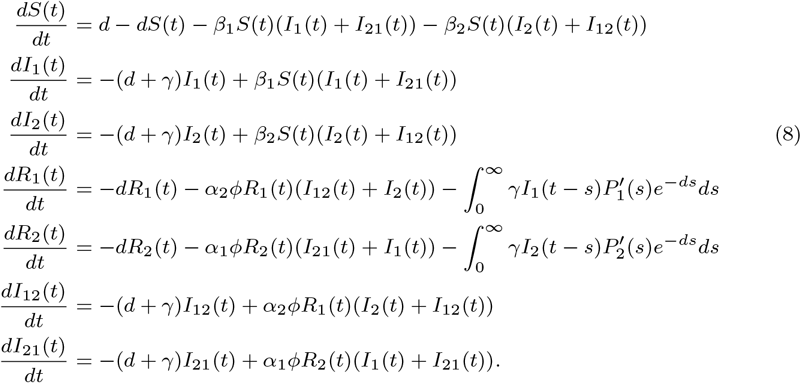

According to Miller (1971), the limiting system ensures that the initial system proposed has an equilibrium. Also, according Hethcote (1981), using Miller’s theorems (Miller, 1971), it is possible to show that the equilibrium of the system coincide with those of their limiting equation.

For this limiting system (8) it is necessary to define the Banach space with memory, as defined by Feng (2017), Wang (2012), Li (2010) and Röst (2008), in order to have a well-posed system and, to have solutions defined in (−∞, 0]. For qualitative theory will be useful to consider the following space phase.

Denote *Δ*_1_, *Δ*_2_ positives constants such that *Δ*_*i*_ < *d*, satisfying, for *i* = 1, 2, 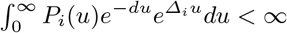. Define the Banach space, for *i* = 1, 2,

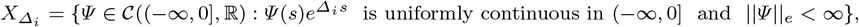

where 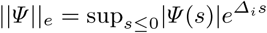. We consider 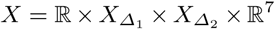 as the phase space for the limiting system (8).

Denote 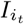, for *i* = 1, 2, the solution *I*_*i*_(*t*) at time *t*, that is 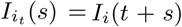, *s* ≤ 0. Let 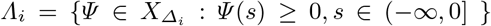, *i* = 1, 2. Then, for initial conditions, 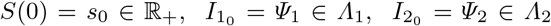, *C*_*i*_(0) = *c*_*i*_ ∈ ℝ_+_, *R*_*i*_(0) = *r*_*i*_ ∈ ℝ_+_, *I*_*ji*_(0) = *θ*_*i*_ ∈ ℝ_+_, *i* = 1, 2, the solutions of the limiting systems in *X* remain non-negative and, 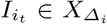, for all *t*, for *i* = 1, 2. Moreover, 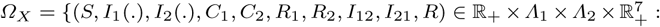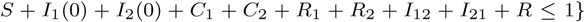 is positively invariant for system (8).

The trivial equilibrium *D*_0_ = (1, 0, 0, 0, 0, 0, 0, 0, 0, 0) always is in the invariant set *Ω*_*X*_. Now using the assumption

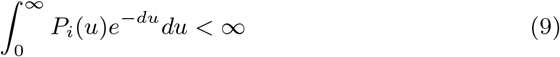

we get 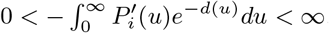. Thus, we will define

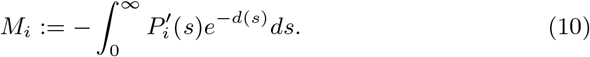

Furthermore, we also assume that

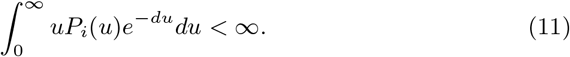

The assumption (11) together with (9) leads to

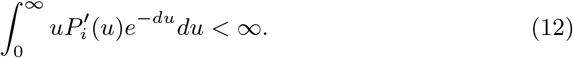

Since 0 < *M*_*i*_ < 1, in the case of the extinction of one of the strains, the BE of the system (8) are

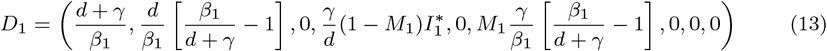

and,

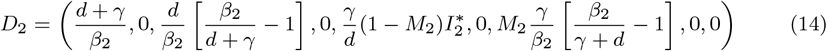

and it will be in the *Ω*_*X*_ region, as long as the parameters satisfy 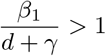 and, 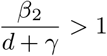, respectively.

In the case of coexistence of the two strain, the CEE is given by

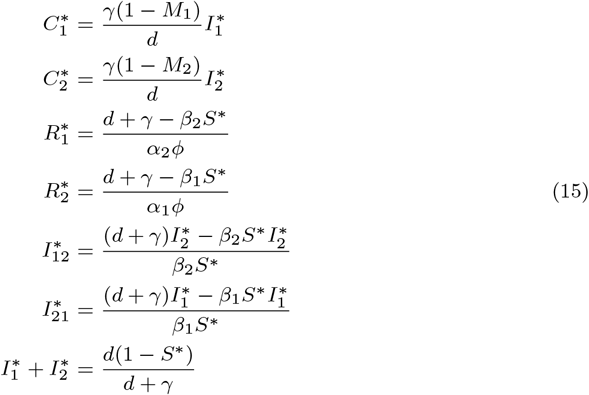

and, *S*^*\**^ is the root of the cubic polynomial *O*(*S*) = *b*_3_*S*^3^ + *b*_2_*S*^2^ + *b*_1_*S* + *b*_0_ where

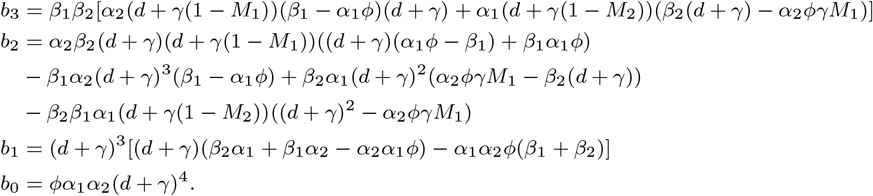

Moreover, *S*^*\**^ has to satisfies that 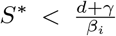, *i* = 1, 2, in order to have the equilibrium in *Ω*_*X*_. This discussion prove the following theorems about the equilibriums of the system (8).

### Theorem 1

*If* 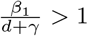 *then the system of equations (8), always has a BE, D*_1_, *in Ω*_*X*_. *And, if* 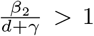 *then the system of equations (8), always has a BE, D*_2_, *in Ω*_*X*_.

### Theorem 2

*Without loss of generality, assume β*_2_ > *β*_1_. *If* 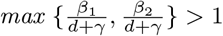 *and*,

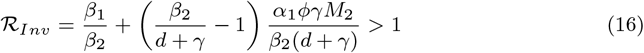

*then, the system (8) admits an equilibrium of the coexistence with two strains (a CEE)*.

*Proof* The independent term, *b*_0_, of the polynomial *O*(*S*) is always positive. Since the equilibrium is given by (15) with *S*^*\**^ being a root of the polynomial *O*, and this equilibrium will be in the region *Ω* if 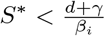, for *i* = 1, 2 let us define

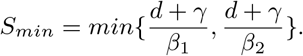

Then, if 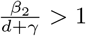 and ℛ_*Inv*_ > 1 then

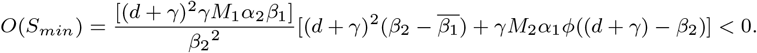

This shows that we have a root *S*^*\**^ of the polynomial *O*, such that, 0 < *S*^*\**^ < *S*_*min*_ for *i* = 1, 2. Therefore, for this *S*^*\**^, we have a positive equilibrium in *Ω*_*X*_ with the coexistence of the two strains. □

### 3.1 Stability Analysis

In section (3) we calculated the equilibriums of the limiting system in order to know the equilibriums of the delay system. In this section, we are going to introduced results that connect the local stability of the limiting system with the local stability of the delay system.

For the purpose of finding the stability of the solutions of the system (5) we follow the study conducted by Brauer (1978), denoting the limiting system (8) as the unperturbed system.

So, we regard

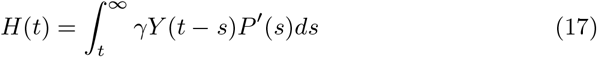

as a perturbation, where *H* is a vector that contains functions, *Y* is the matrix containing the variables of the populations, *P*′ is the vector containing the functions 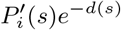. This perturbation function tends explicit to zero as *t* goes to infinity. Adding the perturbation (17) with the limiting system (8) we have the initial system (5).

The stability of the equilibriums of the limiting system is a consequence of stability of the zero solution of the linearized system. And, the asymptotic stability of the zero solution of the linear system 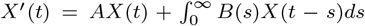 is equivalent to find no solutions in the right half plane *Reλ* ≥ 0 of 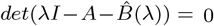, where *I* is the identity matrix, *A* is a matrix and 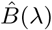 denote the Laplace transform of *B* (Grossman, 1973).

Since, the assumptions (12) hold and, the perturbation function of the system is integrable, we have the necessary assumptions for use the Theorem 2 in Brauer (1978). First, we need the following results about the stability of the limiting system (8), defining mathematically 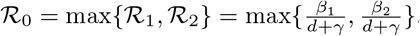.

#### Theorem 3

*If* ℛ_0_ = *max*{ℛ_1_, ℛ_2_} < 1 *then the DFE (D*_0_*), of the system (8) is locally asymptotically stable. And, D*_0_ *will be unstable if* ℛ_0_ > 1.

*Proof* Solving the characteristic equation

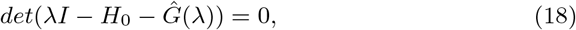

where *I* is the identity matrix 9 × 9, *H*_0_ is the matrix

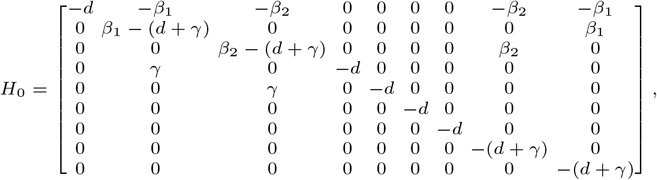

and,

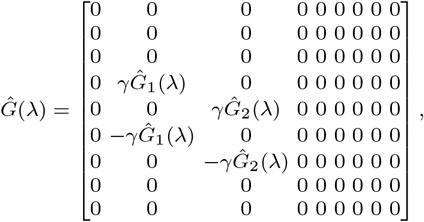

With 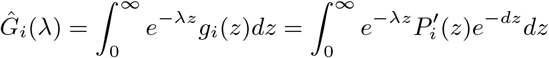, gives us the following eigenvalues of the system at DFE:

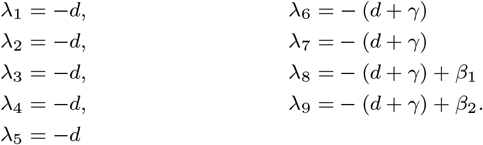

If ℛ_0_ < 1 then all the eigenvalues are negative. If ℛ_0_ > 1 then at least one eigenvalue will be positive. This guarantees the prove. □

*Remark 1* Without loss of generality, we are assuming that the infection has different forces and, therefore, we assume for now on that *β*_2_ > *β*_1_.

#### Theorem 4

*The BE, D*_1_, *of the system (8) is always unstable in Ω*_*X*_ *region. And, the BE, D*_2_, *is locally stable, in Ω*_*X*_, *if* ℛ_*Inv*_ < 1 *and, is unstable in Ω*_*X*_, *if* ℛ_*Inv*_ > 1.

*Proof* Solving the characteristic equation

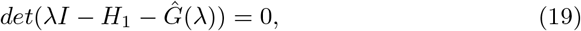

where *I* is the identity matrix 9 × 9, *H*_1_ is the matrix

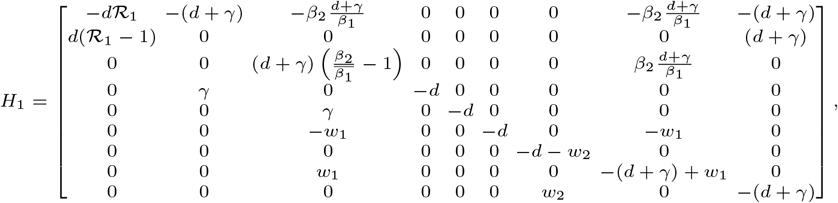

Where 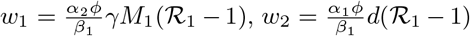 and, 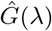 is the matrix whose the elements are the Laplace transform of the 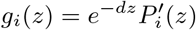, of the associated linear system, gives us the following eigenvalues of the system at *D*_1_, given by (13):

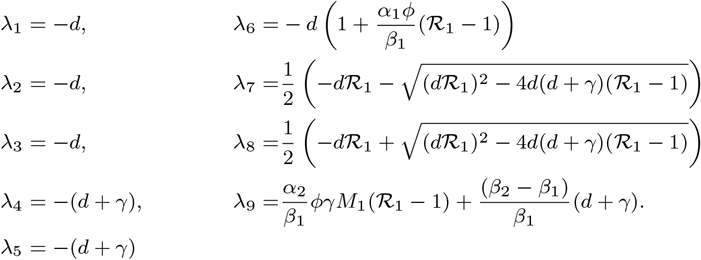

Since *β*_2_ > *β*_1_, the eigenvalue *λ*_9_ will be always positive. While, solving the characteristic equation of the associated linear system in *D*_2_, given by (14), gives us the following eigenvalues:

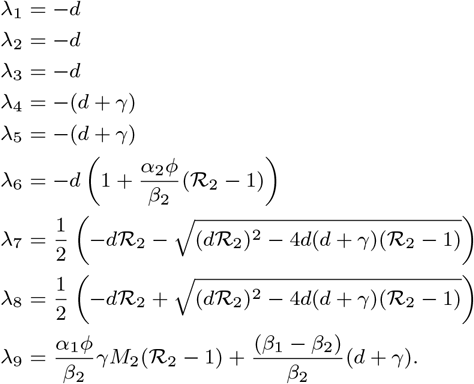

Therefore, all the eigenvalues has negative real part, except for one eigenvalue that change its sign. If ℛ_*Inv*_ < 1 the eigenvalue *λ*_9_ is negative. In the other hand, if ℛ_*Inv*_ > 1 thus, *λ*_9_ is positive. This prove the result. □

*Remark 2* Since *M*_2_ < 1 then, if 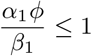, we have that ℛ _*Inv*_ < ℛ_1_. Biologically speaking this results means that there is a range of values for *β*_1_ for which the strain one can not invade the population if the strain two is endemic. In this way, the infection forces two may protect the population from infection forces one. After this range value, the strains coexist.

#### 3.1.1 Stability of the solutions of the Time Delay System

We showed at section (3) the equilibriums of the unperturbed system (8) and the theorems proved uniformly asymptotic stability of the zero solution of the linear limiting system, and therefore, the stability of the equilibriums of the limiting system. Thus, using the Theorem 2 (Brauer, 1978) we have the following results about the stability of solutions of the initial system (5) and, therefore for (4).

##### Corollary 1

*If* ℛ_0_ < 1, *thus the DFE, D*_0_, *of the system (5) is locally asymptotically stable. And D*_0_ *is unstable if* ℛ_0_ > 1.

##### Corollary 2

*The BE, D*_1_, *of the system (5) is always unstable in Ω. And, the BE, D*_2_, *is locally stable in Ω if* ℛ_*Inv*_ < 1 *and, is unstable in Ω if* ℛ_*Inv*_ > 1.

*Remark 3* The analysis of the local stability of the CEE using this theory was not successful, since we have to deal with a characteristic transcendental equation, with infinite many roots. Thus, we will study the stability of the CEE numerically at section (5).

## 4 Global Stability

In this section, we will investigate the global dynamics of the system (5), by constructing suitable Lyapunov function.

### 4.1 Global Stability of the DFE

#### Theorem 5

*If* ℛ_0_ ≤ 1 *then the DFE, D*_0_, *of the system (5) is globally attractive in Ω*.

*Proof* First, denote the positive function, for *i* = 1, 2,

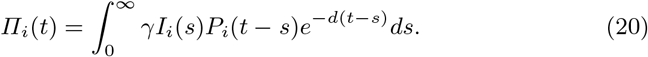

Consider 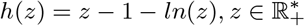. Then, *h* 0. Let *L* be the function

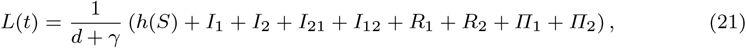

so,

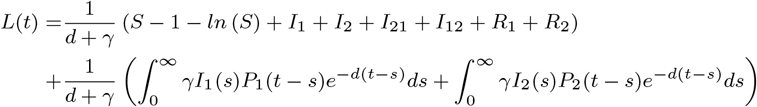

is well-posed, is a Lyapunov function, with *L* ≥ 0, where the equality being true if and only if *S* = 1 and, *I*_*i*_ = 0, *R*_*i*_ = 0, *I*_*ij*_ = 0 for *i, j* = 1, 2, since *h* (*S*) ≥ 0, for any *S* > 0.

Now, differentiating *L* along the solution of the system (5) we have

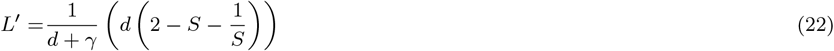

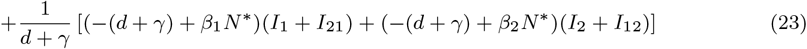

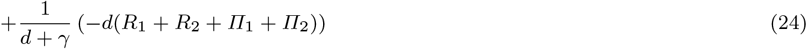

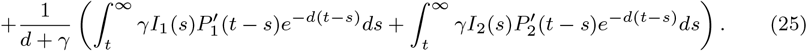

Since 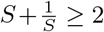 we have that the term in (22) is always non positive in *Ω*. Also, since ℛ_0_ ≤ 1 we have that the term in (23) is always non positive in *Ω. R*_1_ and *R*_2_ are in *Ω* and, *Π*_1_ and *Π*_2_ are positive functions of *t* so we have that the term in (24) its always non positive. The last term in (25) is non positive because *P*_*i*_(*t*) is decreasing and, thus 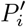 is negative. Therefore, *L* ≤ 0 in *Ω* and, the equality is true if and only if each term of the equation is zero. From (22) the equality is true if and only if *S* = 1 and, with ℛ_0_ < 1, from (23) to (25) we conclude that *I*_1_ = *I*_2_ = *R*_1_ = *R*_2_ = *C*_1_ = *C*_2_ = *I*_12_ = *I*_21_ = 0.

In the case that ℛ_0_ = 1, since *S* = 1, from the first equation of the system we have that *S* = − *β*_*i*_(*I*_*i*_ + *I*_*ji*_) < 0. But this is a contradiction with the fact that, since *S* = 1, implies that *S* = 0. Thus *I*_*i*_ + *I*_*ji*_ = 0. Therefore, for instance, if ℛ_0_ = 1 we still have that *I*_2_ = 0 and *I*_12_ = 0. In that way, defining *E* = {(*S, I*_1_, *I*_2_, *C*_1_, *C*_2_, *R*_1_, *R*_2_, *I*_12_, *I*_21_, *R*) ∈ *Ω* ; *L*^*′*^ (*t*) = 0} thus, the singleton *D*_0_ is the largest invariant set in *E*. By the Invariance Principle for IDE (Burton, 2005; LaSalle, 1976) we have that the DFE, *D*_0_, of the system (5) is globally asymptotically stable in *Ω*. □

### 4.2 Global Stability of the Boundary Equilibrium

We prove, in previously section (3.1.1), that for *β*_2_ > *β*_1_ the BE, *D*_2_, is locally asymptotically stable, when ℛ_2_ > 1 and ℛ_*Inv*_ < 1. Now we are able to show that under this same conditions the trajectories with initials conditions in *Ω* − {(*S, I*_1_, *I*_2_, *C*_1_, *C*_2_, *R*_1_, *R*_2_, *I*_12_, *I*_21_, *R*) ∈ *Ω* ; *I*_2_ = 0} approach that equilibrium. We need, first, the following result.

#### Lemma 1

*If* ℛ_2_ > 1 *and* ℛ_*Inv*_ < 1 *then the trajectories of system (5) that start in Ω approach the invariant set Ω*_2_ = {(*S, I*_1_, *I*_2_, *C*_1_, *C*_2_, *R*_1_, *R*_2_, *I*_12_, *I*_21_, *R*) ∈ *Ω* ; *I*_1_ = *C*_1_ = *R*_1_ = *I*_12_ = *I*_21_ = *R* = 0}.

*Proof* Consider the Lyapunov function

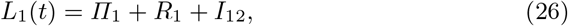

with *Π*_1_ being the same function definite in (20).

Thus, differentiating *L*_1_ along the solution of the system (5) we have

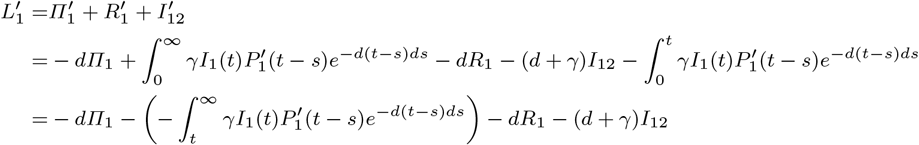

Therefore, 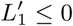 in *Ω* and, the equality is true if, and only if, *R*_1_ = 0, *I*_12_ = 0, *Π*_1_ = 0 and 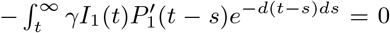. In addition, from the equation of the system and, from the equality above, we have directly that *I*_1_ = 0 and, *C*_1_ goes to zero when t goes to infinity. Also, since *I*_1_ = 0 we have that

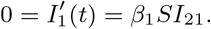

Then, *S* = 0 or *I*_21_ = 0. If *S* = 0 then, for the first equation of the system, we have *S* (*t*) = *d* > 0, which is a contradiction, since *S* = 0 implies *S*^*′*^= 0. Then *I*_21_ = 0. Therefore, we have shown that the maximal invariant set contained in set of all points in *Ω* where 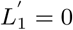 is *Ω*_2_. This shows the lemma. □

This lemma shows that under the some conditions about the Reproduction numbers is sufficient to study the dynamics of the delay system (5) only on the projection of *Ω*_2_, this is, in the set 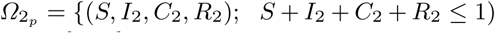 in ℝ^4^. In this set, the initial system is reduced to

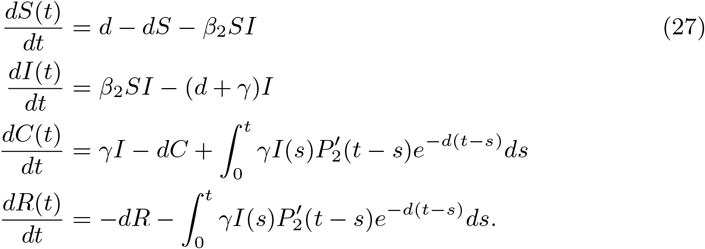

This system have two equilibriums 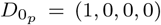 and 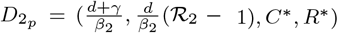 correspondents to the projections of *D*_0_ and *D*_2_, respectively. The following theorem show that all solutions of the system (27) with initial conditions in 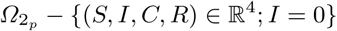 approach the equilibrium *D*_2_ when ℛ_2_ > 1.

#### Theorem 6

*If* ℛ_2_ > 1 *then the Endemic equilibrium* 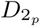 *of the system (27) is globally stable in* 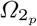.

*Proof* Proof is straightforward from classical SIR Model. Since the first two equations do not depend of C and R, the solutions when, ℛ_0_ > 1, converge to the endemic equilibrium. And, once *I* converges to *I*^*\**^, *C* converges to *C*^*\**^. □

The local asymptotic stability of *D*_2_, the lemma and, the theorem above, demonstrate the global stability of *D*_2_ in *Ω* under the conditions ℛ_2_ > 1 and ℛ_*Inv*_ < 1.

## 5 Numerical Results

In this chapter we are going to use numerical approach to obtain information about the local stability of the CEE. The numerical values for the parameters are shown in the table 1.

**Table 1:**
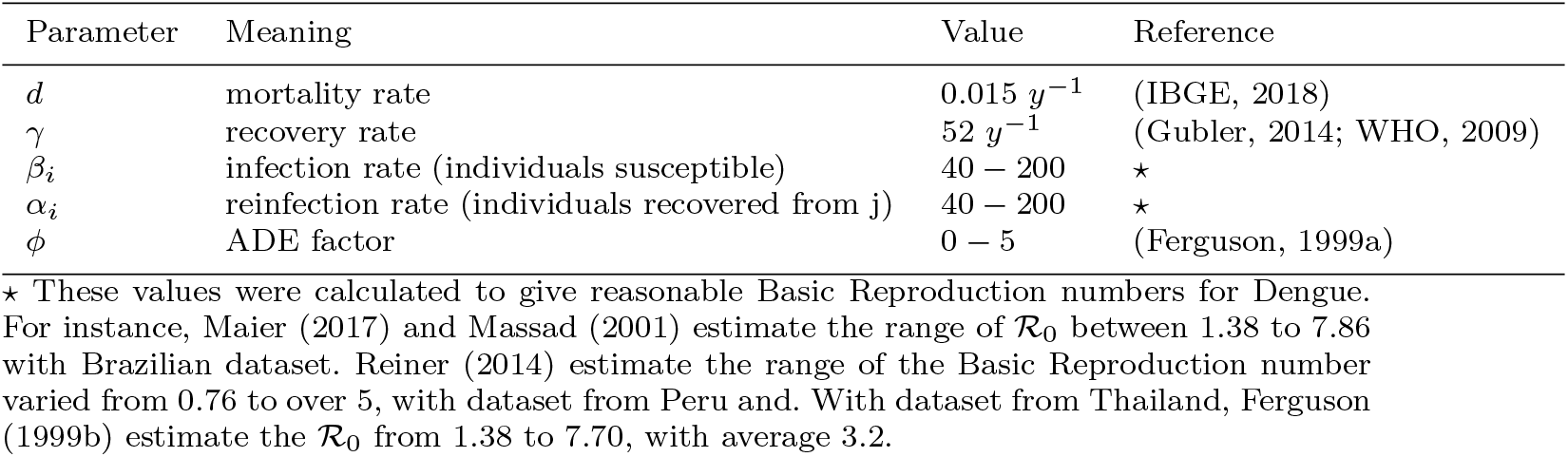
Numerical values of the parameters used in simulations

Is also necessary to choose a function that can represent the immunity period. For this initial numerical analysis we choose to work with the cubic polynomial (in order to not work with infinity time),

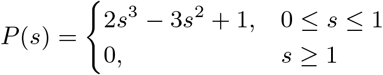

with average time being 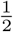 year (Gubler, 2014). In addition, we are assuming that after one year the cross immunity is not effective anymore and, after this period the individual is susceptible again for other strains (Gubler, 2014).

### 5.1 Stability of the Coexistence Endemic equilibrium

As we saw at section (3) the CEE *D*_3_ exists only if ℛ_0_ > 1 and, if ℛ_*Inv*_ > 1. In this case, the BE losses stability and the CEE, with the coexistence of the two strains, rises and remain in the invariant region.

Figures (1a) and (1b) show the parameters region for the stability of the BE and, for the DFE and, the parameters region for the existence of the CEE. Note that, as the value *ϕ* increases, the parameters region of the stability for the BE decreases, forcing the CEE coexist within the region. Additionally, there is a threshold for the value of *ϕ*, in each case, that satisfies ℛ_*Inv*_ > 1.

**Fig. 1:**
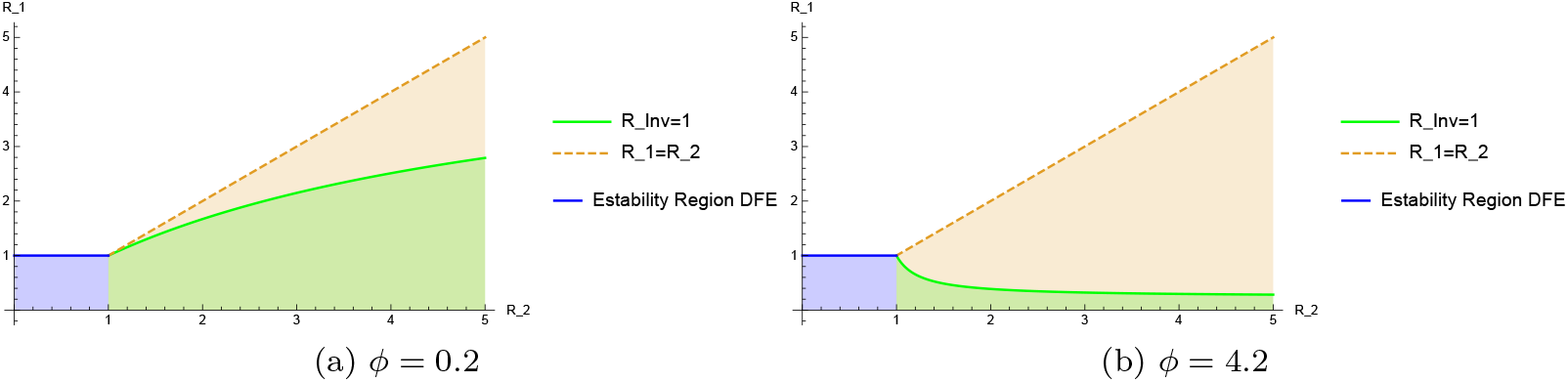
Stability and existence region of the equilibriums. The blue region represents the parameters region for which the DFE is globally stable (ℛ_0_ ≤ 1). The green region represents where the BE is locally stable (ℛ_*Inv*_ ≤ 1). The coral one represents the existence region of the CEE (ℛ_*Inv*_ > 1).

The characteristic equation of the system is a transcendental equation, with typically having infinitely many roots. Thus, we are going to do a numerical analysis in order to study the stability of the CEE, analyzing the roots of the characteristic equation, numerically, for some values of the parameter *ϕ*, using it as a bifurcation parameter.

We separated in two cases:

**Case (i):** ℛ_0_ > 1, ℛ_1_ < 1

In this case, with values for the parameters in table (1) chosen *β*_1_ = 45 and *β*_2_ = 180, which gives ℛ_1_ = 0.87, and ℛ_0_ = 3.46. The ℛ_*Inv*_ is bigger than one for *ϕ* = 1.23. Then, for all value of *ϕ* > 1.23 we have the existence of the CEE, and the correspondents eigenvalues, as show in the figures (2a) to (2f).

**Fig. 2:**
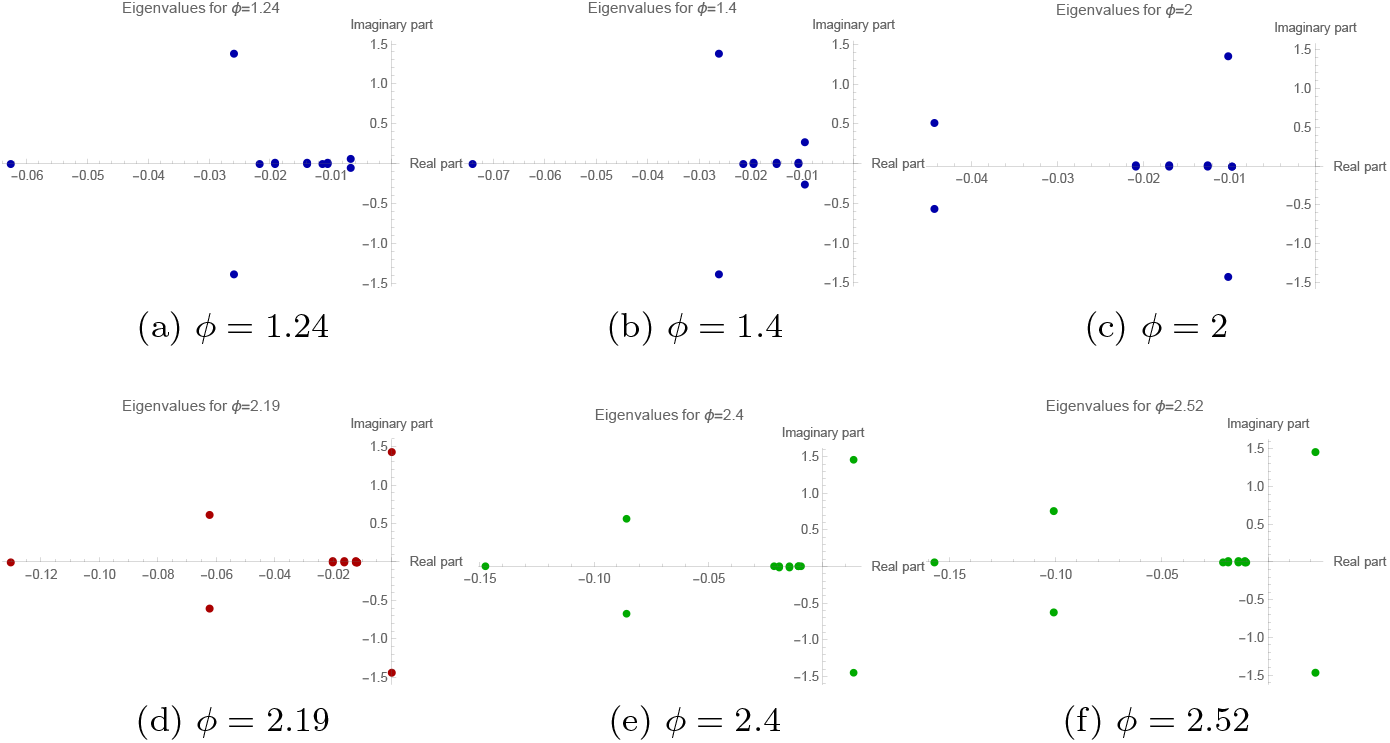
The figures show the eigenvalues of the Endemic equilibrium in the complex plane, for each value of *ϕ*. Figure (2d) shows that a purely imaginary eigenvalue appears for *ϕ* ≈ 2.19.

Figures (2a) to (2f) show that the characteristic equation has a pair of conjugated complex roots, that change the sign of the real part as *ϕ* increases. Thus, a Hopf bifurcation occurs when the parameter *ϕ* ≈ 2.19.

**Case (ii):** ℛ_0_ > 1, ℛ_1_ > 1

In this case, with values for the parameters in table (1) chosen *β*_1_ = 120 and *β*_2_ = 180, which gives ℛ_1_ = 2.31, and ℛ_0_ = 3.46. The ℛ_*Inv*_ is bigger than one for *ϕ* = 0.205. Then, for all value of *ϕ* > 0.205 we have the existence of the CEE, and the correspondents eigenvalues, as show in the figures (3a) to (3f).

**Fig. 3:**
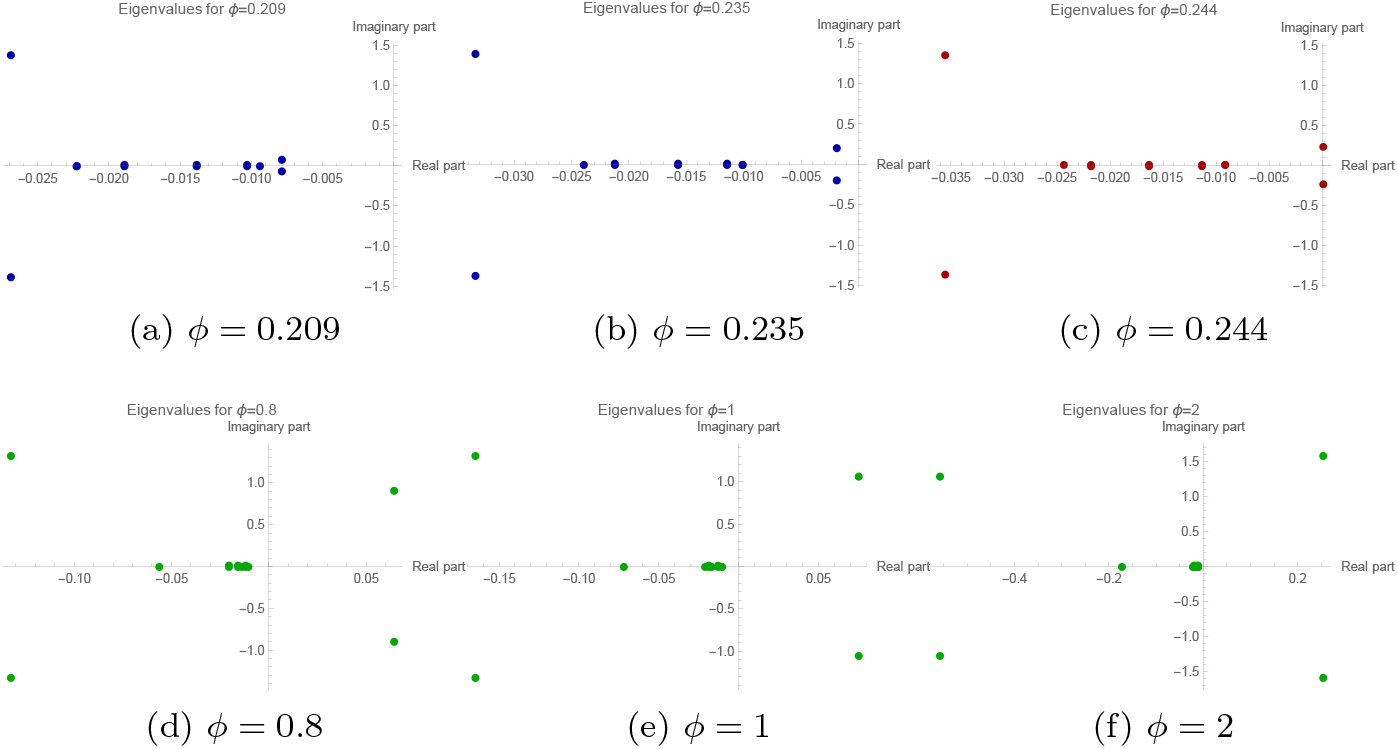
The figures show the eigenvalues of the Endemic equilibrium in the complex plane, for each value of *ϕ*. Figure (3c) show that a purely imaginary eigenvalue appears for *ϕ* ≈ 0.244.

Figures (3a) to (3f) show that the characteristic equation has a pair of conjugated complex roots, that change the sign of the real part as *ϕ* increases. Thus, a Hopf bifurcation occurs when the parameter *ϕ* ≈ 0.244.

Furthermore, after the Hopf Bifurcation, the CEE will be unstable, leading to complex dynamic where sometimes, the infection one resists leading to the other be almost extinct and sometimes, the opposite. Models that carry these mechanisms that can cause the coexistence of pathogens and, therefore, this switches between the strains has an important role, since according to VinodKumar (2013), different serotypes have been coo-circulating in the same area with one of them being dominant during an outbreak.

#### 5.1.1 Bifurcation Structure

As seen in the figures at section (5.1), the CEE change the stability as the parameter *ϕ* changes. As *ϕ* increase from small values trough critical value, *ϕ*_*c*_, the steady state changes from a stable focus to an unstable steady state. Therefore, Hopf bifurcation occurs and thus, we conclude that closed periodic orbit will be found in a small neighbourhood of *ϕ*_*c*_.

The Hopf bifurcation occurs at *ϕ*_*c*_ = 2.19 (figure (4a)) and at *ϕ*_*c*_ = 0.244 (figure (4b)) and, therefore, the solutions exhibit a small amplitude limit cycle around the endemic equilibrium. A stable limit cycle arises close to the critical Bifurcation point and goes away from the unstable equilibrium. Thus, it is possible to conclude that a supercritical Hopf bifurcation occurred.

**Fig. 4:**
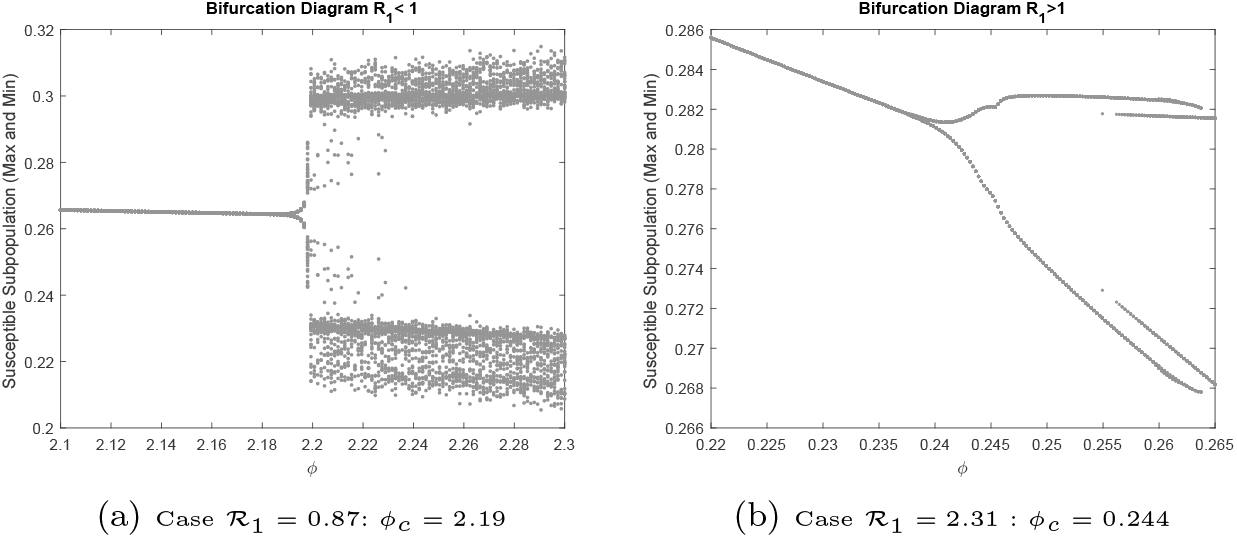
In the horizontal axis, the parameter *ϕ* varies in a vicinity of *ϕ*_*c*_, while in the vertical axis, the maximum and minimum value for susceptible population are plotted.

This change of stability, and thus, the Hopf bifurcation is local. Therefore, the Hopf bifurcation does not specify what happens when the parameter is further beyond the vicinity of its critical bifurcation value Edelstein-Keshet (2005); Murray (2002). Because of that, solutions will be plotted, in the next section, for different values of *ϕ*, to show the asymptotic behaviour also for parameter values further from bifurcation value.

#### 5.1.2 Solutions of the system

**Case (i):** ℛ_0_ > 1, ℛ_1_ < 1

Figures (5a) to (5e) show the solutions of the system for the case ℛ_1_ < 1 and ℛ_0_ > 1. Figure (5a) for *ϕ* = 0.5 and ℛ_*Inv*_ < 1. Thus, as ℛ_0_ > 1 the solution converges to the value of BE.

**Fig. 5:**
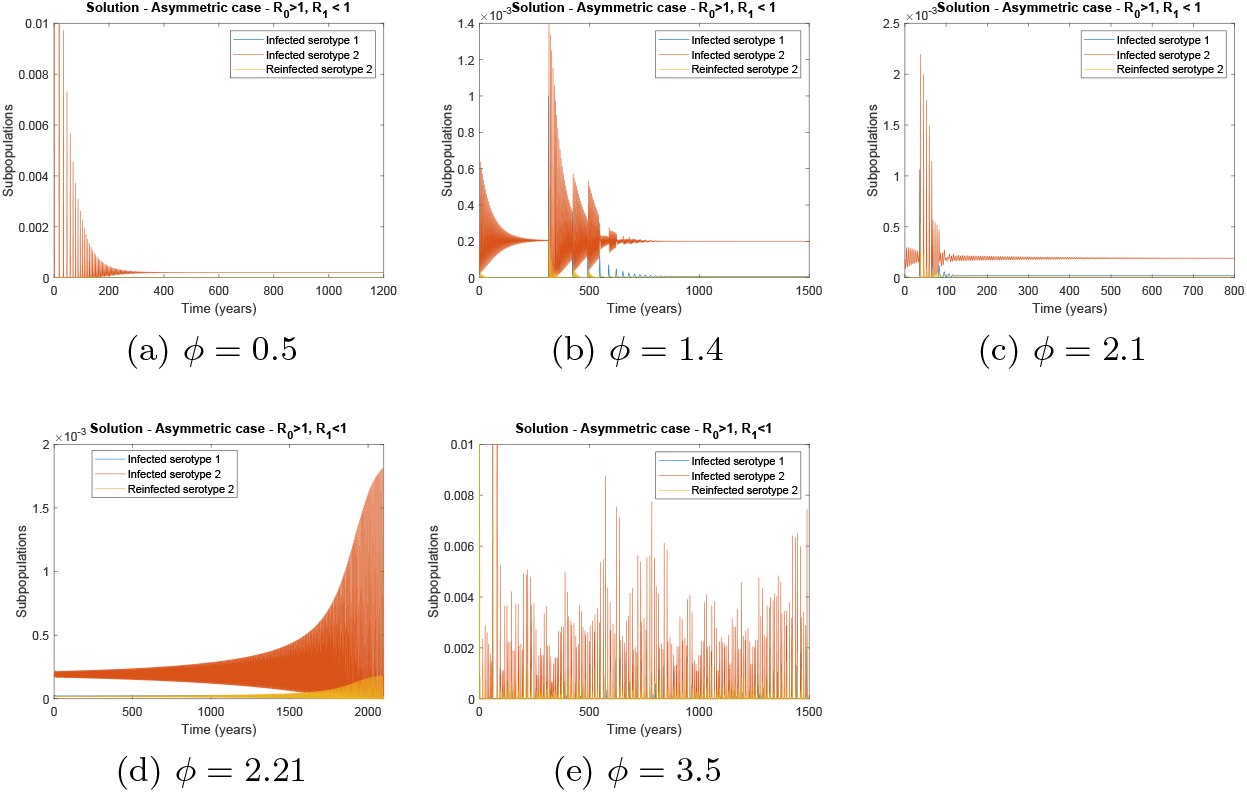
Times series for different values of *ϕ*, in case (i), with ℛ_1_ = 0.87.

Figures (5b) and (5c) for *ϕ* = 1.4 and *ϕ* = 2.1, respectively, where ℛ_*Inv*_ > 1. Since *ϕ* is small than the critical value of HB, thus the solution converges to the CEE, this is, the CEE is stable.

Figure (5d) for *ϕ* = 2.21 and ℛ_*Inv*_ > 1. Since the value of *ϕ* is closed to the HB critical value, periodic solutions is seen. Figure (5e) for *ϕ* = 3.5 where the parameter is bigger than the the HB critical value. Thus, for long term behaviour the solutions seen not to converge to periodic orbit, showing complex dynamic.

**Case (ii):** ℛ_0_ > 1, ℛ_1_ > 1

Figures (6a) to (6e) show the solutions of the system for the case ℛ_1_ > 1 and ℛ_0_ > 1. Figure (6a) for *ϕ* = 0.1 the ℛ_*Inv*_ < 1. Thus, as ℛ_0_ > 1 the solution converges to the value of BE. Figure (6b) for *ϕ* = 0.23, smaller than the HB critical value, thus the solution converges to the CEE, this is, the CEE is stable.

**Fig. 6:**
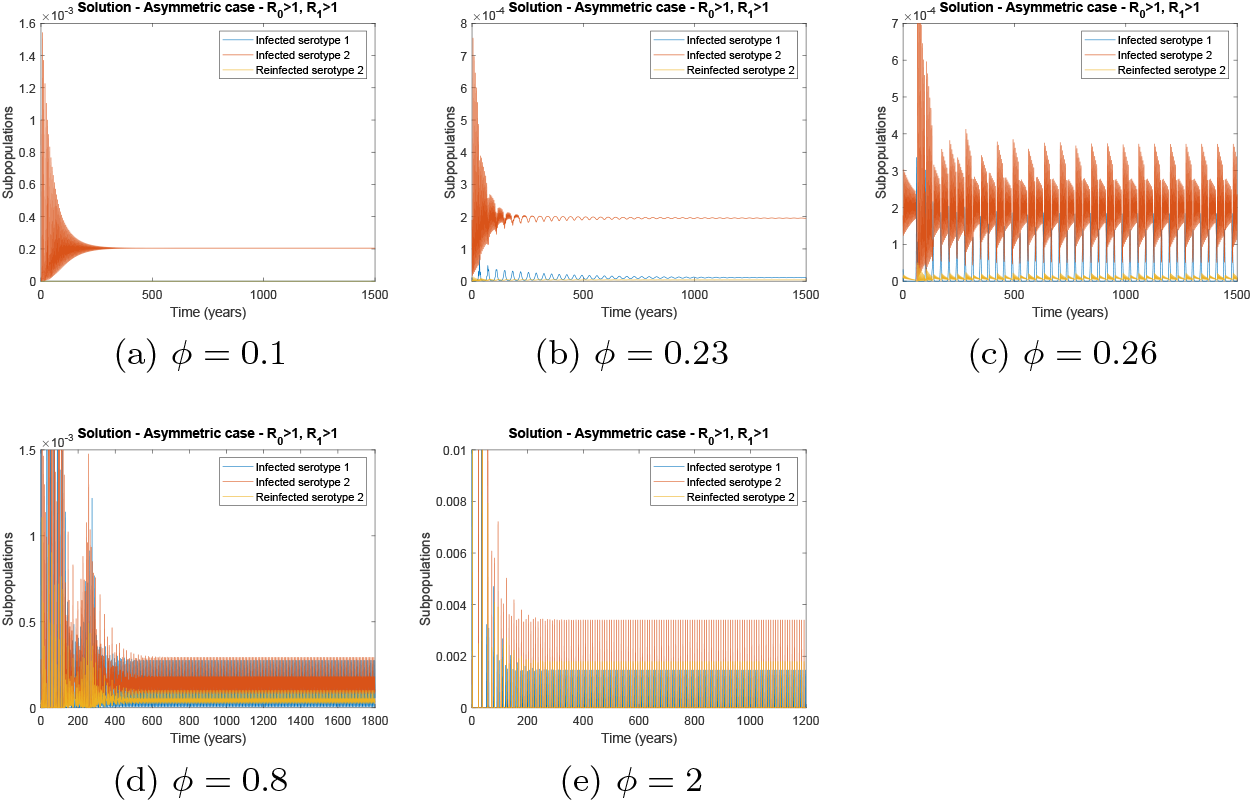
Times series for different values of *ϕ*, in case (ii), with ℛ_1_ = 2.31.

Figure (6c) show the solution for ℛ_*Inv*_ > 1 and *ϕ* = 0.26, closed to the HB critical value, thus periodic solutions is seen. Figure (6d) for *ϕ* = 0.8 and the ℛ_*Inv*_ > 1, where the parameter is bigger than the HB critical value. Thus, for long term behaviour the solutions seen to converge to periodic orbit. Figure (6e) for *ϕ* = 2 bigger than the HB critical value. Also, in this case, solutions converge to periodic orbits.

Although we conclude numerically that the solutions of the system, for values of the parameter far from bifurcation critical value, go to an equilibrium or to a periodic orbit, for some cases, the solutions seem not to converge, showing complex dynamics.

Therefore, biologically speaking, since the value for ADE parameter is unknown it is hard to give a smallest interval that is a representative value for ADE. This means that it is hard to predict the next episode of the disease, since we can have different scenarios: coexistence of infections, periodic outbreaks, or even complex dynamics, depending on parameter interval value for ADE.

## 6 Discussion and Conclusions

In this work we have developed a mathematical model that can be applied to a multi strain infectious disease, as Dengue fever. Dengue fever is endemic in more than 100 countries (WHO, 2018) and remains a major public health problem.

Over the years, some mathematical models for infectious diseases and specially for Dengue fever have been developed, most of which are in the format of ODEs.

However, epidemics propagation are not instantaneous and it is more appropriate to model epidemics incorporating continuously distributed delay and not discrete delay or constant time.

We proposed a model with time delay on temporary immunity, allowing general periods of immunity. The time delay was used to model phenomena that an individual may not be immediately susceptible after being recovered. Also, a constant parameter describe the ADE effect.

A special form for the temporary immunity function was considered in section (2). By considering the temporary immunity time exponential distributed allowed us to reduce the IDE system to an ODE system. Thus, the qualitative analysis of the ODE system gave us a visual picture of the dynamic behaviour.

In the section (3), using results from IDE systems we are capable of to prove the existence of four equilibriums and the stability of the equilibriums, proving that the disease will die out if ℛ_0_ < 1. If the ℛ_*Inv*_ < 1, one strain die out and the other will persist. And, in the last scenario the two strains will coexist if ℛ_*Inv*_ > 1. Lyapunov functions can be an effective tool for proving global stability of IDE systems, although it may be difficult to find it. Nevertheless, we could prove the global stability of DFE and of the BE when ℛ_*Inv*_ < 1. This result means, biologically, that there is a range of values for the infection force for which the strain one can not invade the population if the strain two is endemic. In this way, the infection two may protect the population from infection one.

Comparing the particular case (exponential distributed function) with the general case, the qualitative behaviour of the system was not altered by the distributed delay. However, the Invasion number depends on the average cross protection time, that depends on the choice of the function, which can affect whether the infection could coexist.

The ADE effect was study by considering it as a bifurcation parameter. Thus, the possibility for Hopf bifurcation was examined through a numerical analysis. This kind of bifurcation is local, therefore only for values closed to the bifurcation critical point we can conclude that a limit cycle exists and, therefore, the solutions will show periodic behaviour. It means that, in the scenario that the strains coexist, the strains can coexist in an equilibrium or, for specific value, they can show periodic behaviour, according to the value of the ADE parameter.

For solutions of the system, for values further from bifurcation critical value, numerical results show that the solutions go to an equilibrium or go to a periodic orbit. However, analysis of other kinds of bifurcations will be necessary.

Since ADE is a factor still unknown, we only can give scenarios as periodic solutions or coexistence of strains, as well as, complex dynamics, depending on this value. Although a possible outbreak will be hard to predict.

Lastly, mathematical models focus in understanding the spread of infectious disease as well as suggesting interventions to control or even predicting the consequences of the propagation. With this model we tried to understand the dynamic of different strains at population level, as well as, explain why is so hard to predict Dengue fever. And we concluded that there is some mechanism and intrinsic characteristics of Dengue fever, such as ADE and cross protection, which may hinder the prediction of the next outbreaks of the disease.

## Acknowledgements

This research was supported by Coordenação de Aperfeiçoamento de Pessoal de Nível Superior - Brazil (CAPES) - Finance Code 001 and LIAM – Laboratory for Industrial and Applied Mathematics, Department of Mathematics and Statistics, York University-CA. The author thank Fernando Valdes Ravelo for his help with Matlab program.

