## Supplementary Material for "Cross immunity protection and antibody dependent enhancement: A distributed delay dynamic model"

**Noname manuscript No.**
(will be inserted by the editor)

### Supplementary Material - Cross immunity protection and antibody dependent enhancement: A distributed delay dynamic model

**Vanessa Steindorf · Sergio Oliva · Jianhong Wu**

Received: date / Accepted: date

#### 1 Particular case: Exponential immunity

Here we are going to analyse qualitatively the ODE system, a particular case of the model studied. If we assume the length of immunity being exponentially distributed, which means that the fraction of temporarily immune individual remain in the $C_i$ class is $P^i(t) = e^{-\omega_i t}$, with $\omega_i > 0$, for $i = 1, 2$, then the IDE system became an ODE system. Thereby, the system already normalized, where the variables represent the fractions of the populations, can be describe as follows:

CAPES and LIAM

University of São Paulo, Institute of Mathematics and Statistics, Applied Mathematics Department
Rua do Matão 1010, São Paulo - SP, 05508-090, Brazil


University of São Paulo, Institute of Mathematics and Statistics, Applied Mathematics Department
Rua do Matão 1010, São Paulo - SP, 05508-090, Brazil


York University, Laboratory for Industrial and Applied Mathematics, Department of Mathematics & Statistics, Faculty of Science and Engineering
4700 Keele St., Toronto, Canada, M3J 1P3


$$\begin{aligned}
\frac{dS(t)}{dt} &= d - dS(t) - \beta_1 S(t)(I_1(t) + I_{21}(t)) - \beta_2 S(t)(I_2(t) + I_{12}(t)) \\
\frac{dI_1(t)}{dt} &= -(d+\gamma)I_1(t) + \beta_1 S(t)(I_1(t) + I_{21}(t)) \\
\frac{dI_2(t)}{dt} &= -(d+\gamma)I_2(t) + \beta_2 S(t)(I_2(t) + I_{12}(t)) \\
\frac{dC_1(t)}{dt} &= -dC_1(t) + \gamma I_1(t) - \omega_1 C_1(t) \\
\frac{dC_2(t)}{dt} &= -dC_2(t) + \gamma I_2(t) - \omega_2 C_2(t) \\
\frac{dR_1(t)}{dt} &= -dR_1(t) - \alpha_2 \phi R_1(t)(I_{12}(t) + I_2(t)) + \omega_1 C_1 \\
\frac{dR_2(t)}{dt} &= -dR_2(t) - \alpha_1 \phi R_2(t)(I_{21}(t) + I_1(t)) + \omega_2 C_2 \\
\frac{dI_{12}(t)}{dt} &= -(d+\gamma)I_{12}(t) + \alpha_2 \phi R_1(t)(I_2(t) + I_{12}(t)) \\
\frac{dI_{21}(t)}{dt} &= -(d+\gamma)I_{21}(t) + \alpha_1 \phi R_2(t)(I_1(t) + I_{21}(t)).
\end{aligned} \tag{1}$$

Clearly, the system (1) always has a Disease-free equilibrium (DFE), namely,

$$E_0 = (1, 0, 0, 0, 0, 0, 0, 0, 0, 0).$$

In the case of the extinction of one of the strains we are able to find the Boundary equilibrium (BE) of the system

$$E_1 = \left( \frac{d+\gamma}{\beta_1}, \frac{d}{\beta_1}\left[\frac{\beta_1}{d+\gamma} - 1\right], 0, \frac{\gamma}{d+\omega_1} I_1^*, 0, \frac{\omega_1}{d}\frac{\gamma}{d+\omega_1} I_1^*, 0, 0, 0, 0 \right). \tag{2}$$

And, the BE

$$E_2 = \left( \frac{d+\gamma}{\beta_2}, 0, \frac{d}{\beta_2}\left[\frac{\beta_2}{d+\gamma} - 1\right], 0, \frac{\gamma}{d+\omega_2} I_2^*, 0, \frac{\omega_2}{d}\frac{\gamma}{d+\omega_2} I_2^*, 0, 0, 0 \right). \tag{3}$$

For biological reasons we are looking for steady states that belongs to $\Omega$ a positively invariant region, where $\Omega = \{(S, I_1, I_2, C_1, C_2, R_1, R_2, I_{12}, I_{21}, R) \in \mathbb{R}_+^{10}$ such that $S + I_1 + I_2 + C_1 + C_2 + R_1 + R_2 + I_{12} + I_{21} + R \leq 1\}$. In this way, the BE, $E_i$, $i = 1, 2$ is in $\Omega$ as long as $\dfrac{\beta_i}{d+\gamma} > 1$.

In the case of the coexistence of the two strains, we are able to find the Coexistence Endemic equilibrium (CEE) that is given by

$$\begin{aligned} C_1^* &= \frac{\gamma}{d+\omega_1} I_1^* \\ C_2^* &= \frac{\gamma}{d+\omega_2} I_2^* \\ R_1^* &= \frac{d+\gamma-\beta_2 S^*}{\alpha_2\phi} \\ R_2^* &= \frac{d+\gamma-\beta_1 S^*}{\alpha_1\phi} \\ I_{12}^* &= \frac{(d+\gamma)I_2^* - \beta_2 S^* I_2^*}{\beta_2 S^*} \\ I_{21}^* &= \frac{(d+\gamma)I_1^* - \beta_1 S^* I_1^*}{\beta_1 S^*} \\ I_1^* + I_2^* &= \frac{d(1-S^*)}{d+\gamma} \end{aligned} \tag{4}$$

and, $S^*$ is the root of the cubic polynomial $O(S) = b_3 S^3 + b_2 S^2 + b_1 S + b_0$ where

$$\begin{aligned} b_3 &= \beta_1\beta_2[\alpha_2(\beta_1 - \alpha_1\phi)(d+\gamma)(d+\omega_2)((d+\omega_1)(d+\gamma) - \gamma\omega_1) \\ &\quad + \alpha_1(\beta_2(d+\gamma)(d+\omega_1) - \alpha_2\phi\gamma\omega_1)((d+\omega_2)(d+\gamma) - \gamma\omega_2)] \\ b_2 &= (d+\gamma)^3(d+\omega_1)(d+\omega_2)[\beta_1\alpha_2(-\beta_1+\alpha_1\phi) + \beta_2\alpha_2(-\beta_1+\alpha_1\phi) - \beta_2\alpha_1(\beta_1+\beta_2)] \\ &\quad + (d+\gamma)^2\beta_1\beta_2((d+\omega_2)\gamma\omega_1\alpha_2 + (d+\omega_1)\gamma\omega_2\alpha_1) \\ &\quad + \beta_1\beta_2\alpha_1\alpha_2\phi[(d+\gamma)^2(d+\omega_1)(d+\omega_2) - \omega_2\gamma^2\omega_1] \\ b_1 &= (d+\gamma)^3(d+\omega_1)(d+\omega_2)[(d+\gamma)(\beta_2\alpha_1 + \beta_1\alpha_2 - \alpha_2\alpha_1\phi) - \alpha_1\overline{\alpha_2}\phi(\beta_1+\beta_2)] \\ b_0 &= \phi\alpha_1\alpha_2(d+\gamma)^4(d+\omega_1)(d+\omega_2). \end{aligned}$$

Moreover, $S^*$ has to satisfies that $S^* < \frac{d+\gamma}{\beta_i}$, $i = 1,2$, in order to have the CEE, $E_3$, in the $\Omega$ region. Otherwise, if $S^*$ does not satisfies this inequality, the variables that represents recovered and infected population will be negative.

This discussions support and demonstrate the following theorems.

**Theorem 1** *If* $\frac{\beta_1}{d+\gamma} > 1$ *then the system of equations (1), always has a BE,* $E_1$*, given by (2) in* $\Omega$*. And, if* $\frac{\beta_2}{d+\gamma} > 1$ *then the system of equations (1), always has a BE,* $E_2$*, given by (3) in* $\Omega$*.*

**Theorem 2** *Without loss of generality, suppose that* $\beta_2 > \beta_1$*. If* $max\{\frac{\beta_1}{d+\gamma}, \frac{\beta_2}{d+\gamma}\} > 1$ *and,*

$$\mathcal{R}_{Inv} = \frac{\beta_1}{\beta_2} + \left(\frac{\beta_2}{d+\gamma} - 1\right)\frac{\alpha_1\phi\gamma\omega_2}{\beta_2(d+\omega_2)(d+\gamma)} > 1 \tag{5}$$

*then, the system (1) admits a CEE in* $\Omega$ *with the coexistence of the two strains.*

*Proof* The independent term, $b_0$, of the polynomial $O(S)$ is always positive. Since the equilibrium is given by (4) with $S^*$ being a root of the polynomial $O$ and, this equilibrium will be in the region $\Omega$ if $S^* < \frac{d+\gamma}{\beta_i}$, for $i = 1,2$ let us define

$$S_{min} = \min\{\frac{d+\gamma}{\beta_1}, \frac{d+\gamma}{\beta_2}\} = \frac{d+\gamma}{\beta_2}.$$

Then, if $\frac{\beta_2}{d+\gamma} > 1$ and $\mathcal{R}_{Inv} > 1$, the polynomial $O$ at $S_{min}$ is

$$O(S_{min}) = \frac{[(d+\gamma)^2\gamma\omega_1\alpha_2\beta_1]}{\beta_2^2}[(d+\omega_2)(d+\gamma)^2(\beta_2-\beta_1)+\gamma\omega_2\alpha_1\phi((d+\gamma)-\beta_2)] < 0.$$

This shows that we have a root $S^*$ of the polynomial $O$, such that, $0 < S^* < S_{min}$. Therefore, for this $S^*$ we have a positive equilibrium of the system (1), in $\Omega$, with the coexistence of the two strains. □

#### 1.1 Basic Reproduction number

We defined the Basic Reproduction number in the subsection 2.1.1 of this study. We defined

$$\mathcal{R}_1 = \frac{\beta_1}{d+\gamma} \tag{6}$$

as the Basic Reproduction n for strain one. And,

$$\mathcal{R}_2 = \frac{\beta_2}{d+\gamma} \tag{7}$$

as the Basic Reproduction number for strain two.

As well as the overall Reproduction number for the system was defined as $\mathcal{R}_0 = \max\{\mathcal{R}_1, \mathcal{R}_2\}$.

Another important threshold value, the Invasion Reproduction number, was defined in that subsection, and it is given by

$$\mathcal{R}_{Inv} = \frac{\mathcal{R}_1}{\mathcal{R}_2} + (\mathcal{R}_2 - 1)\frac{\overline{\alpha_1}\phi\gamma\omega_2}{\overline{\beta_2}(d+\omega_2)(d+\gamma)}. \tag{8}$$

*Remark 1* For our purpose and without loss of generality, we are assuming that the infection has different forces and, therefore, we assume for now on that $\beta_2 > \beta_1$. Thus $\mathcal{R}_0 = \mathcal{R}_2$.

#### 1.2 Stability Analysis

The local stability of the equilibriums will be determined by the classical method of determining stability of the steady states of some ODE system, by analysis of the eigenvalues of the Jacobian matrix of the system at each equilibrium.

**Theorem 3** *If* $\mathcal{R}_0$ *then the DFE,* $E_0$*, of the system (1) is locally asymptotically stable. And,* $E_0$ *is unstable if* $\mathcal{R}_0 > 1$.

*Proof* The eigenvalues of Jacobian matrix of the system evaluated at the DFE ($E_0$) are

$$\begin{aligned}
\lambda_1 &= -d\\
\lambda_2 &= -(d+\omega_1)\\
\lambda_3 &= -(d+\omega_2)\\
\lambda_4 &= -(d+\gamma)\\
\lambda_5 &= -(d+\gamma)\\
\lambda_6 &= -d\\
\lambda_7 &= -d\\
\lambda_8 &= -(d+\gamma)+\beta_1\\
\lambda_9 &= -(d+\gamma)+\beta_2.
\end{aligned}$$

Clearly if $\mathcal{R}_0 < 1$ thus all the eigenvalues will be negative. If $\mathcal{R}_0 > 1$ thus at least one eigenvalue will be positive. Which proves the theorem. □

**Theorem 4** *The BE, $E_1$, of the system (1) is always unstable, in $\Omega$ region. And, the BE, $E_2$, is stable in $\Omega$, if $\mathcal{R}_{Inv} < 1$ and, unstable in $\Omega$, if $\mathcal{R}_{Inv} > 1$.*

*Proof* The eigenvalues of the Jacobian matrix of the system evaluated at $E_1$ are

$$\begin{aligned}
\lambda_1 &= -d\\
\lambda_2 &= -(d+\omega_1)\\
\lambda_3 &= -(d+\omega_2)\\
\lambda_4 &= -(d+\gamma)\\
\lambda_5 &= -(d+\gamma)\\
\lambda_6 &= -d\left(1+\frac{\alpha_1\phi}{\beta_1}(\mathcal{R}_1-1)\right)\\
\lambda_7 &= \frac{1}{2}\left(-d\mathcal{R}_1-\sqrt{(d\mathcal{R}_1)^2-4d(d+\gamma)(\mathcal{R}_1-1)}\right)\\
\lambda_8 &= \frac{1}{2}\left(-d\mathcal{R}_1+\sqrt{(d\mathcal{R}_1)^2-4d(d+\gamma)(\mathcal{R}_1-1)}\right)\\
\lambda_9 &= \frac{\alpha_2\phi\gamma\omega_1(\mathcal{R}_1-1)}{\beta_1(d+\omega_1)}+\frac{(\beta_2-\beta_1)(d+\gamma)(d+\omega_1)}{\beta_1(d+\omega_1)}.
\end{aligned}$$

Since $\beta_2 > \beta_1$, the eigenvalue $\lambda_9$ will be always positive. In addition, the eigenvalues of the Jacobian matrix of the system evaluated at $E_2$ are

$$\begin{aligned}
\lambda_1 &= -d\\
\lambda_2 &= -(d+\omega_1)\\
\lambda_3 &= -(d+\omega_2)\\
\lambda_4 &= -(d+\gamma)\\
\lambda_5 &= -(d+\gamma)\\
\lambda_6 &= -d\left(1+\frac{\alpha_2\phi}{\beta_2}(\mathcal{R}_2-1)\right)\\
\lambda_7 &= \frac{1}{2}\left(-d\mathcal{R}_2-\sqrt{(d\mathcal{R}_2)^2-4d(d+\gamma)(\mathcal{R}_2-1)}\right)\\
\lambda_8 &= \frac{1}{2}\left(-d\mathcal{R}_2+\sqrt{(d\mathcal{R}_2)^2-4d(d+\gamma)(\mathcal{R}_2-1)}\right)\\
\lambda_9 &= \frac{\alpha_1\phi\gamma\omega_2(\mathcal{R}_2-1)}{\beta_2(d+\omega_2)}+\frac{(\beta_1-\beta_2)(d+\gamma)(d+\omega_2)}{\beta_2(d+\omega_2)}.
\end{aligned}$$

If $\mathcal{R}_{Inv} < 1$ thus, the eigenvalue $\lambda_9 < 0$. Furthermore, the others eigenvalue has negative real part. Therefore, the BE, $E_2$, will be stable.

If $\mathcal{R}_{Inv} > 1$ thus, the eigenvalue $\lambda_9 > 0$. This proves the theorem. □

*Remark 2* We can rewrite the threshold $\mathcal{R}_{Inv}$ as

$$\mathcal{R}_{Inv} = \frac{\mathcal{R}_1}{\mathcal{R}_2} + \frac{\alpha_1\phi}{\beta_1}\frac{\gamma}{(d+\gamma)}\frac{\omega_2}{(d+\omega_2)}\mathcal{R}_1\left(1-\frac{1}{\mathcal{R}_2}\right). \tag{9}$$

Thus, if $\dfrac{\alpha_1\phi}{\beta_1} \leq 1$, we have that $\mathcal{R}_{Inv} < \mathcal{R}_1$. This results means that there is a range of values for $\beta_1$ for which the strain one can not invade the population if the strain two is endemic. In this way, the strain two may protect the population from strain one.

The analysis of the local stability of the CEE using the classical theory was not successful. We will have to deal with a characteristic polynomial of a $9 \times 9$ matrix and, with the fact that it was not possible to describe the value of $S^*$ in terms of the parameters. Thus, in order to study the stability we will analyse it numerically, at section (2).

#### 2 Numerical Analysis

The numerical values for the parameters are shown in the table 1.

##### 2.1 Stability of the Coexistence Endemic equilibrium

In this section, we are going to explore the stability of the CEE numerically. As we saw at section (1) the CEE, $E_3$, exists only if $\mathcal{R}_0 > 1$ and $\mathcal{R}_{Inv} > 1$. In this case, the BE losses stability and the CEE, with the coexistence of the two strains, rises and remain in the invariant region.

Figures (1a) and (1b) show the parameters region for the stability of the BE and, for the DFE and, the parameters region for the existence of the CEE. Note

Table 1: Numerical values of the parameters used in simulations

| Parameter | Meaning | Value | Reference |
|---|---|---|---|
| $d$ | mortality rate | $0.015\ y^{-1}$ | (IBGE, 2018) |
| $\gamma$ | recovery rate | $52\ y^{-1}$ | (Gubler, 2014; WHO, 2009) |
| $\omega_i$ | cross immunity protection rate | $2\ y^{-1}$ | (Gubler, 2014; WHO, 2009) |
| $\beta_i$ | infection rate (individuals susceptible) | $40 - 200$ | $\star$ |
| $\alpha_i$ | reinfection rate (individuals recovered from j) | $40 - 200$ | $\star$ |
| $\phi$ | ADE factor | $0 - 5$ | (Ferguson, 1999a) |

$\star$ These values were calculated to give reasonable Basic Reproduction numbers for Dengue. For instance,Maier (2017) and Massad (2001) estimate the range of $\mathcal{R}_0$ beteween 1.38 to 7.86 with Brazilian dataset. Reiner (2014) estimate the range of the Basic Reproduction number varied from 0.76 to over 5, with dataset from Peru and. With dataset from Thailand, Ferguson (1999b) estimate the $\mathcal{R}_0$ from 1.38 to 7.70, with average 3.2.

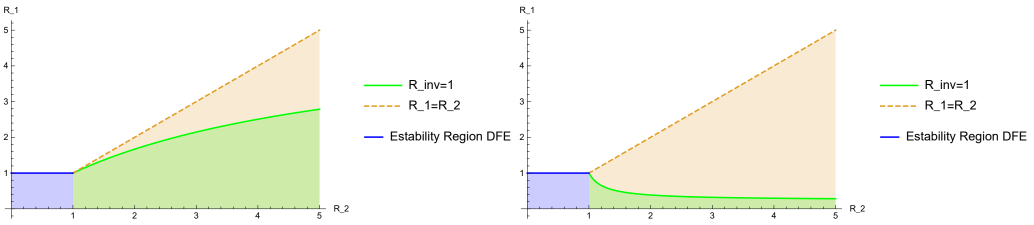


(a) $\phi = 0.2$ (b) $\phi = 4.2$

Fig. 1: Stability and existence region of the equilibriums. The blue region represents the parameters region for which the DFE is globally stable ($\mathcal{R}_0 \leq 1$). The green region represents where the BE is locally stable ($\mathcal{R}_{Inv} \leq 1$). The coral one represents the existence region of the CEE ($\mathcal{R}_{Inv} > 1$).

that, as the value $\phi$ increases, the parameters region of the stability for the BE decreases, forcing the CEE coexist within the region.

To show the stability of the CEE numerically we need to guarantee that the values of the parameters satisfy $\mathcal{R}_{Inv} > 1$. $\phi$, the parameter used to describe the ADE effect, is a parameter that its value is unknown. As we saw in figures (1a) and (1b) there is a threshold for the value of $\phi$, in each case, that satisfies $\mathcal{R}_{Inv} > 1$. Starting for this threshold value, the CEE will be in the positive region and, therefore, we can look for the eigenvalues of the Jacobian Matrix at CEE. Note that the CEE can be in the invariant region even if the strain one is not established. Thus, we separated in two cases:

**Case (i):** $\mathcal{R}_0 > 1$**,** $\mathcal{R}_1 < 1$

In this case, with values for the parameters in table (1), chosen $\beta_1 = 45$ and $\beta_2 = 180$, which gives $\mathcal{R}_1 = 0.87$, and $\mathcal{R}_0 = 3.46$. The $\mathcal{R}_{Inv}$ is bigger than one for $\phi$ bigger than 1.23. Then, for all value of $\phi > 1.23$ we have the CEE and, the correspondents eigenvalues, as show in the figures (2a) to (2f).

Figures (2a) to (2f) give the stability of the CEE, $E_3$, for each value of $\phi$. We can see that the matrix has two conjugated complex roots that change the sign of the real part as $\phi$ increases. Thus, a Hopf bifurcation occurs when the parameters $\phi \approx 2.52$ (figure (2d)).

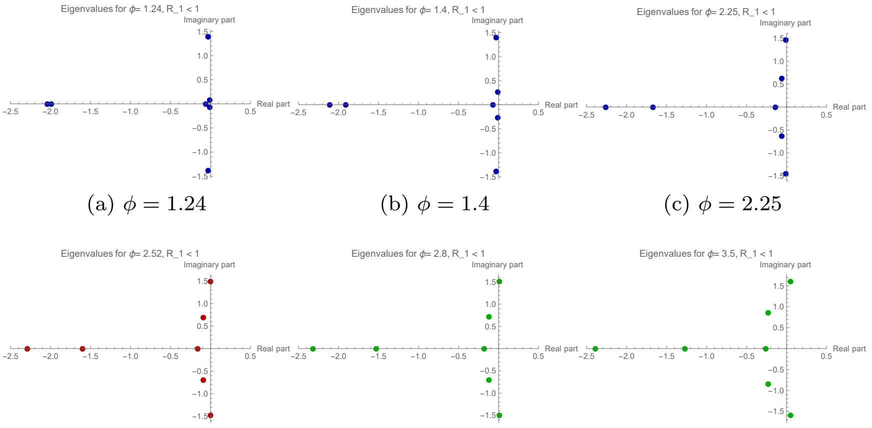


(a) $\phi = 1.24$ (b) $\phi = 1.4$ (c) $\phi = 2.25$

(d) $\phi = 2.52$ (e) $\phi = 2.8$ (f) $\phi = 3.5$

Fig. 2: Eigenvalues of the Endemic equilibrium in the complex plane, for each value of $\phi$. Figures (2a) to (2f) show that the real part of a pair of complex eigenvalues change the sign as $\phi$ increases.

**Case (ii):** $\mathcal{R}_0 > 1$**,** $\mathcal{R}_1 > 1$

In this case, with the values for the parameters in the table (1) chosen $\beta_1 = 120$ and $\beta_2 = 180$, which gives $\mathcal{R}_1 = 2.31$, and $\mathcal{R}_0 = 3.46$. The $\mathcal{R}_{Inv}$ is bigger than one for $\phi$ bigger than 0.21. Then, for all value of $\phi > 0.21$ we have the CEE and, the correspondents eigenvalues, as show in the figures (3a) to (3f).

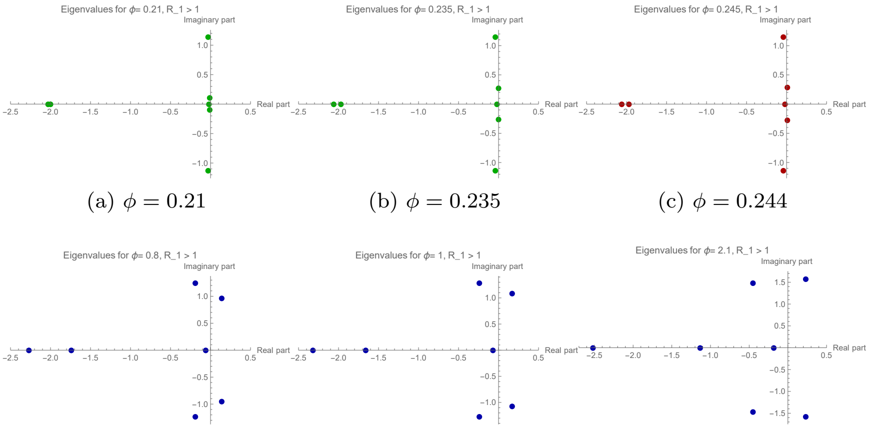


(a) $\phi = 0.21$ (b) $\phi = 0.235$ (c) $\phi = 0.244$

(d) $\phi = 0.8$ (e) $\phi = 1$ (f) $\phi = 2.1$

Fig. 3: Eigenvalues of the Endemic equilibrium in the complex plane, for each value of $\phi$. Figures (3a) to (3f) show that the real part of a pair of complex eigenvalues change the sign as $\phi$ increases.

Figures (3a) to (3f) give the stability of the CEE, $E_3$, for each value of $\phi$. We can see that the matrix has two conjugated complex roots, that change the sign of the real part as $\phi$ increases. Thus, a Hopf bifurcation occurs when parameters $\phi \approx 0.244$ (figure (3c)).

##### *2.1.1 Bifurcation Structure*

As seen in the figures at section (2.1), as $\phi$ increase from small values trough critical value, $\phi_c$, the steady state changes from a stable focus to an unstable steady state. Therefore, Hopf bifurcation occurs and thus, we conclude that closed periodic orbit will be found in a small neighbourhood of $\phi_c$. In order to see the limit cycle around the equilibrium, in a small vicinity of the critical value, a bifurcation diagram are shown in the figures (4a) and (4b).

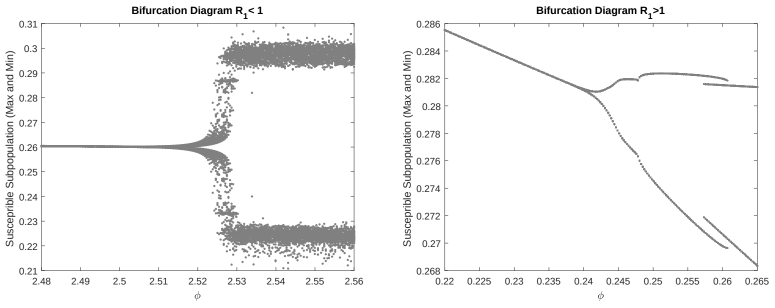


(a) Case $\mathcal{R}_1 = 0.87$: $\phi_c = 2.52$

(b) Case $\mathcal{R}_1 = 2.31$: $\phi_c = 0.244$

Fig. 4: In the horizontal axis, the parameter $\phi$ varies in a vicinity of $\phi_c$, while in the vertical axis, the maximum and minimum value for susceptible population are plotted.

The Hopf bifurcation occurs at $\phi_c = 2.52$ (figure (4a)) and at $\phi_c = 0.244$ (figure (4b)) and, therefore, the solutions exhibit a small amplitude limit cycle around the CEE. A stable limit cycle arises close to the critical Bifurcation point and goes away from the unstable equilibrium. Thus, it is possible to conclude that a supercritical Hopf bifurcation occurred.

This change of stability, and thus, the Hopf bifurcation is local. Therefore, the Hopf bifurcation does not specify what happens when the parameter is further beyond the vicinity of its critical bifurcation value (Edelstein-Keshet, 2005; Lynch, 2004; Murray, 2002). Because of that, solutions will be plotted, in the next section, for different values of $\phi$, to show the asymptotic behaviour also for parameter values further from bifurcation value.

#### 2.2 Solutions of the system: Particular case

In this section we are going to explore numerically the solutions of the system in order to obtain information about the dynamic also for values of $\phi$ further from the bifurcation value.

**Case (i):** $\mathcal{R}_0 > 1$, $\mathcal{R}_1 < 1$

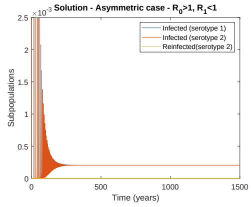


(a) $\phi = 0.5$

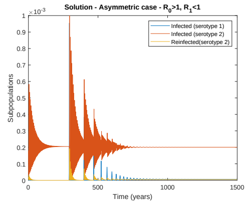


(b) $\phi = 1.4$

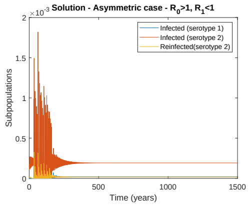


(c) $\phi = 2.1$

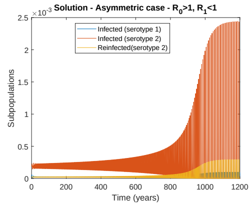


(d) $\phi = 2.55$

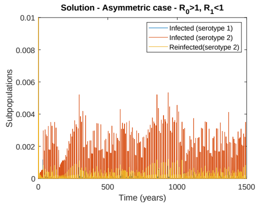


(e) $\phi = 3.5$

Fig. 5: Times series for different values of $\phi$, in case (i), with $\mathcal{R}_1 = 0.87$.

Figures (5a) to (5e) show the solutions of the system for the case $\mathcal{R}_1 < 1$ and $\mathcal{R}_0 > 1$. Figure (5a) for $\phi = 0.5$ and $\mathcal{R}_{Inv} < 1$. Thus, as $\mathcal{R}_0 > 1$ the solution converges to the value of BE.

Figures (5b) and (5c) the $\mathcal{R}_{Inv} > 1$, for $\phi = 1.4$ and $\phi = 2.1$ respectively. Since the value of $\phi$ is smaller than the value of $\phi_c$, thus the solutions converge to the CEE, this is, the CEE is stable. Figure (5d) the $\mathcal{R}_{Inv} > 1$, for $\phi = 2.55$ close to $\phi_c$. Thus, periodic solutions is seen.

Figure (5e) the $\mathcal{R}_{Inv} > 1$, for $\phi = 3.5$ bigger than the value of $\phi_c$. For long term behaviour the solutions seen not to converge to periodic orbit, showing complex dynamic.

**Case (ii):** $\mathcal{R}_0 > 1$, $\mathcal{R}_1 > 1$

Figures (6a) to (6e) show the solutions of the system for the case $\mathcal{R}_1 > 1$ and $\mathcal{R}_0 > 1$. Figure (6a) for $\phi = 0.1$ the $\mathcal{R}_{Inv} < 1$. Thus, as $\mathcal{R}_0 > 1$ the solution converges to the value of BE.

Figure (6b) the $\mathcal{R}_{Inv} > 1$, for $\phi = 0.23$, smaller than the value of $\phi_c$, thus the solution converges to the CEE, this is, the CEE is stable. Figure (6c) the $\mathcal{R}_{Inv} > 1$, for $\phi = 0.26$ close to $\phi_c$. Thus, periodic solutions is seen.

Figure (6d) the $\mathcal{R}_{Inv} > 1$, for $\phi = 0.8$ bigger than the value of $\phi_c$. For long term behaviour the solutions seen to converge to periodic orbit. Figure (6e) the

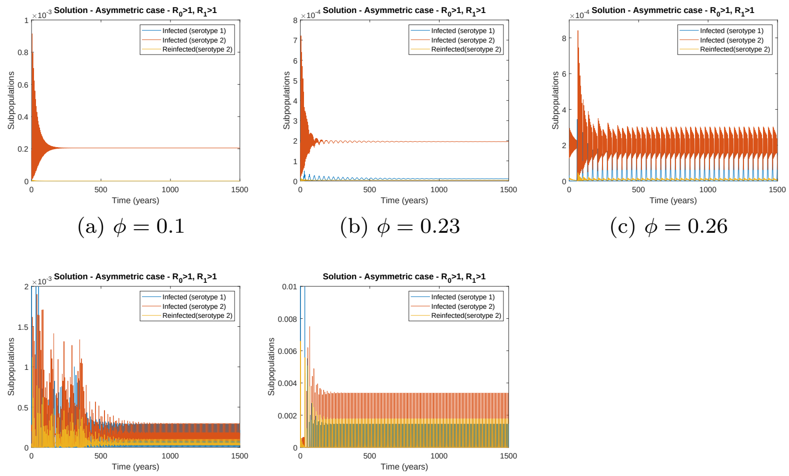


(a) $\phi = 0.1$ (b) $\phi = 0.23$ (c) $\phi = 0.26$

(d) $\phi = 0.8$ (e) $\phi = 2$

Fig. 6: Times series for different values of $\phi$, in case (ii), with $\mathcal{R}_1 = 2.31$.

$\mathcal{R}_{Inv} > 1$, for $\phi = 2$ bigger than the value of $\phi_c$. Also, in this case, solutions converge to periodic orbits.

Although we conclude numerically that the solutions of the system go to an equilibrium or to a periodic orbit, for values further from bifurcation value, in some cases, solutions do not converge, showing complex dynamics. Thus, analysis of other kinds of bifurcations would be necessary.

Therefore, biologically speaking, since the value for ADE parameter is unknown it is hard to give a smallest interval that is a representative value for ADE. This means that it is hard to predict the next episode of the disease, since we can have different scenarios: coexistence of infections, periodic outbreaks, or even complex dynamics, depending on parameter interval value for ADE.
